# Antero-posterior gradients of cell plasticity and proliferation modulate posterior regeneration in the annelid *Platynereis*

**DOI:** 10.1101/2023.06.12.544593

**Authors:** Loïc Bideau, Loeiza Baduel, Marianne Basso, Pascale Gilardi-Hebenstreit, Vanessa Ribes, Michel Vervoort, Eve Gazave

## Abstract

Regenerative abilities are extremely variable among animals and may be substantial in some phyla, such as the annelids. So far, the cellular mechanisms underlying regeneration in annelids remain elusive. To precisely determine the origin(s), plasticity and fate of the cells participating in the blastema formation during posterior regeneration in the annelid *Platynereis dumerilii*, we developed specific tools to track proliferative cells as well as gut epithelial cells. We showed that two populations of progenitors are at play during regeneration and that, among them, gut progenitors from differentiated tissues are lineage-restricted. Strikingly, gut progenitors from less differentiated and more proliferative tissues are much more plastic and can produce ectodermal and mesodermal derivatives, in addition to gut cells. However, their plasticity is *de facto* limited as exemplified by their inability to regenerate populations of stem cells responsible for the constant growth of the worms. We evidenced that those stem cells are from local origin (*i.e.* from the segment abutting the amputation plan) as most of the blastema cells. Our results are in favour of a hybrid and flexible cellular model for posterior regeneration in *Platynereis* relying on a gradient of cell plasticity along the antero-posterior axis of the animal.

## Introduction

Regeneration, defined as the ability to reform a lost body part upon injury, is an tremendous and essential process in animals. Its importance is illustrated by its wide deployment in metazoans, although the extent of tissues that can be regenerated is highly variable from one species to another (Bely and Nyberg, 2010). While mammals can regenerate at best an organ, many other species can perform “extensive” regeneration such as the reformation of a limb (*e.g.* salamanders), a large amputated part of their body axis (*e.g.* annelids) or even their whole body from a small fragment of tissue (*e.g.* cnidarians, planarians), (Bideau et al., 2021). Despite this diversity, three common steps for all regenerative processes can be described: (i) the formation of a wound epithelium enclosing the area of the injury, (ii) the recruitment of progenitors at the wound site which often form a blastema (*i.e.* a mass of proliferative undifferentiated mesenchymal cells covered by an epithelium) and (iii) the growth of the blastema through cell proliferation and its differentiation during a morphogenesis step (Galliot and Ghila, 2010; Tiozzo and Copley, 2015).

Uncovering the origin and fate of the cells participating in the blastema formation has been one of the greatest challenges in the field of regenerative biology for decades(Tanaka and Reddien, 2011). Data from major regeneration models revealed that the blastema can be formed by the activation of stem/progenitor cells whose progeny will populate the blastema, as exemplified by the planarian *Schmidtea mediterranea,* whose regeneration is sustained by adult stem cells called neoblasts (Wenemoser and Reddien, 2010). As pluripotent cells (at least part of them), neoblasts participate to the formation of all missing tissues (Wagner et al., 2011). Alternatively, blastema can be fuelled by post-mitotic cells that dedifferentiate and re-enter cell cycle upon injury, like in urodele limb regeneration (Stocum and Cameron, 2011). In this regenerative process, various local tissues close to the wound dedifferentiate into strictly lineage-restricted progenitors, unable to switch to another lineage during regeneration(Flowers et al., 2017; Kragl et al., 2009).

As such, two opposite models have been broadly defined: the first involves very highly plastic cells that migrate to the wound, the second involves local tissues that dedifferentiate to constitute a pool of diverse progenitors with low plasticity. However, the mechanisms of regeneration are often more complex and in many species and contexts, both dedifferentiated cells as well as tissue-specific resident stem cells participate in the blastema formation (*e.g.* axolotl limb regeneration, (Lin et al., 2021; Sandoval-Guzmán et al., 2014)). Therefore, regeneration may also rely on intermediate models in which the sources of blastema cells can be multiple: stem/progenitor cells and dedifferentiation of mature tissues; and in which the plasticity of the cells participating in the regeneration can vary (Sandoval-Guzmán et al., 2014).

Precise data unravelling the source of blastema cells during regeneration are coming from scarce key regenerative model species but are lacking for the majority of regeneration-competent lineages. An interesting phylum to study regeneration is the annelids that have great, yet diverse regenerative capacities. The majority of annelids can indeed regenerate their posterior and/or anterior parts upon amputation (Özpolat and Bely, 2016). Those regeneration processes have been studied from a long time and in various species (reviewed in (Bely, 2006)) but the precise cellular mechanisms involved in the formation of the blastema remain unclear. The annelid *Platynereis dumerilii* is nowadays emerging as a strong and relevant model to address fundamental regeneration questions (Schenkelaars and Gazave, 2021; Özpolat et al., 2021). This marine worm has the ability, after embryonic development, to grow continuously through a process of posterior elongation that relies on putative stem/progenitor cells localized in a subterminal growth zone (Gazave et al., 2013). Importantly, *Platynereis* has the astounding ability, upon posterior amputation, to regenerate its complex posterior part, *via* the formation of a blastema and through stereotyped steps that we have previously defined (Planques et al., 2019). Upon amputation, the wound heals in 24 h (stage 1 at 1 day post amputation or 1dpa). One day later (stage 2, 2dpa), a small blastema has formed and from then on it will intensely grow. Between stages 2 and 3 (3 dpa), the blastema cells have started to differentiate into various tissues (*e.g.* muscles, nervous system…). At stage 3, the growth zone and the pygidium, which is the posterior-most part of the worm body, have begun to reform. At stage 5 (5 dpa), a new fully functional growth zone has been re-established and allows posterior elongation to resume (Planques et al., 2019).

The cellular and molecular mechanisms controlling this regeneration process are still largely unresolved. In our first study on this topic, we had shown, through S-phase cell labeling coupled with proliferation inhibition experiments, that cell proliferation is absolutely necessary from stage 2 onwards for regeneration to be properly achieved (Planques et al., 2019). Besides, pulse-chase experiments suggested that most of the blastema cells have a local origin, from the segment abutting the amputation plan (Planques et al., 2019). It remains to precisely determine the origin(s), plasticity and fate of the cells participating in the regeneration blastema. To this end, in this study, we developed tools to track proliferative cells as well as gut epithelial cells. We determined that some gut cells from differentiated tissues constitute a population of progenitors uniquely participating in the regeneration of gut epithelial cells. Strikingly, we also showed that upon posteriorization (*i.e.* by giving a posterior identity to previously anterior tissues), those lineage-restricted gut progenitors harbour a higher plasticity as they are able to produce ecto/mesodermal derivatives through regeneration. However, they do not participate in the regeneration of the growth zone stem cells, which are from local origin (*i.e.* from the segment abutting the amputation). Those results argue for the existence of a gradient of plasticity, through regeneration, of intestinal progenitors along the antero-posterior axis of the animal.

## Results

### Cells proliferating during continuous growth contribute to posterior regeneration

Previous results showed that cells proliferating in a context of continuous growth of the worms may participate in the posterior regeneration in *Platynereis* (Planques et al., 2019). To go further into the role of those cells during regeneration, we first analysed the pattern of S-phase cells in *Platynereis* non-amputated juvenile worms with EdU labellings. We exposed uninjured worms to EdU, either for 5 or 48h, to discriminate quickly-cycling cells (Planques et al., 2019) from cells harbouring a slower replication rate, which have been shown to contribute to regeneration in several species *e.g.* mammals (Karmakar et al., 2020; Koren et al., 2022) or planarians (Molinaro et al., 2021). The distributions of EdU+ cells were analysed immediately after these exposures both on whole samples (Fig. 1A-B’) and on histological sections (Fig. 1C-D’). Three segments were examined, including the most posterior distinguishable segment (called Segment 1 or S1) (Fig. 1A’, B’, C’’ and D’’) as well as the sixth and seventh segments counted from the posterior region (called Segments 6 and 7 or S6 and S7, respectively) (Fig. 1A, B, C-D’). We showed that the proportion of proliferative cells varied significantly along the antero-posterior body axis. Indeed, there are more EdU+ cells in S1 than in S6 or S7 regardless the EdU incubation time (Fig. 1A, A’, B, B’ and E). Next, we looked at the localization of EdU+ cells within the tissues by performing semi-thin sections of S6 and/or S7 (transversal and longitudinal sections) and S1 (longitudinal sections). Cells labelled with a 5h-long EdU pulse in S6/S7 were mainly delineating a central and circular epithelium, which appears to be the gut (Fig. 1C, C’) (Žídek et al., 2018; Dahlitz et al., 2023).

**Figure 1:**
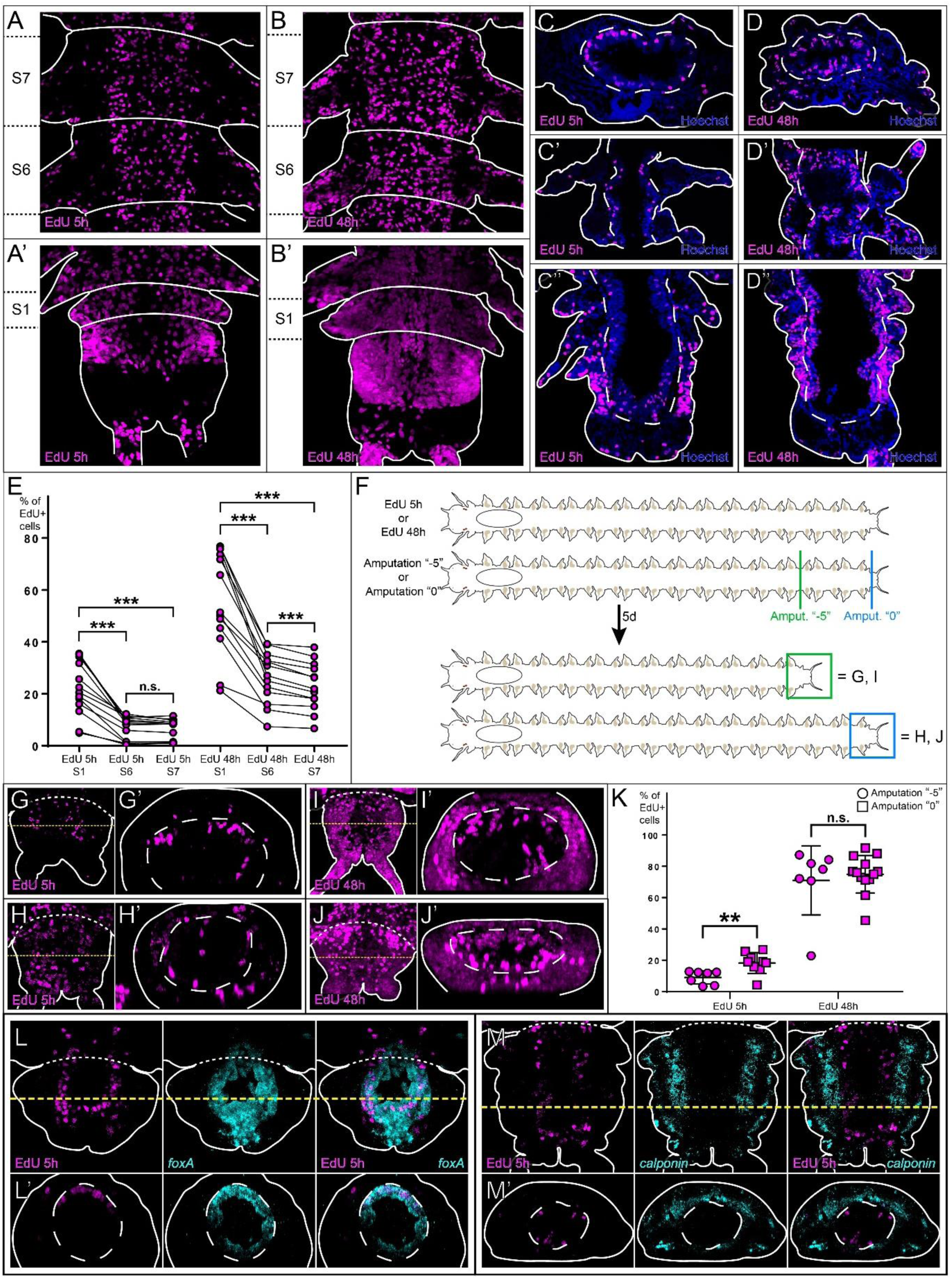
Localization of cells in S-phase in *Platynereis* worms and their contribution to posterior regeneration. **(A-D)** Distribution of EdU+ cells (magenta) and Hoechst DNA staining (blue) in the indicated segments of unamputated juvenile worms after 5h or 48h of EdU incorporation (5µM). A to B’ are confocal z-stacks (ventral views) while C, D are transversal cross-sections and C’ to D’’ longitudinal ones. Anterior is up. EdU+ cells appear to be located mostly inside the gut in C and C’. They are localized in all types of tissues in C’’, D, D’ and D’’. **(E)** Quantification of the proportions of EdU+ cells in the indicated segments and after the specified EdU pulses. Black lines connect values for the same individual. Each bracket corresponds to a Wilcoxon signed-rank test. n.s. p >0.05; *** p <0.001. EdU+ cell proportions are significantly higher in S1 than in S6 and S7 for both EdU pulses. **(F)** Schematic representation of the experiment displayed in G to M. Unamputated juvenile worms were incubated with EdU for 5h or 48h, then amputated at two different positions: amputation “−5” (removal of the pygidium and five segments, depicted in green) or amputation “0” (removal of only the pygidium and the growth zone, depicted in blue). The worms were then let to regenerate for 5 days. **(G-J)** Confocal z-stacks of regenerative parts at stage 5 after either a 5h-long EdU pulse and an amputation “−5” (G) or “0” (H); or after a 48h EdU pulse and an amputation “−5” (I) or “0” (J) (dorsal views, anterior is up). Corresponding virtual transverse sections (along the yellow dotted lines) are shown in G’, H’, I’ and J’ respectively. Dorsal is up. **(K)** Quantification of the proportions of EdU+ cells after the specified EdU pulses and amputation type (circle = amputation “−5”, square = amputation “0”). Each bracket corresponds to a Mann-Whitney U test. n.s. p >0.05; ** p<0.01. Mean + SD are shown. EdU+ cell proportions are higher in H than in G. **(L-M)** Confocal z-stacks of regenerative parts at stage 5 showing EdU+ cells (after a 5h-long EdU pulse prior to an amputation “−5”, magenta) as well as *in situ* hybridization signal (cyan) for gut (*foxA,* L) and smooth muscle markers (*calponin,* M). Single and combined labelling are displayed, all images are dorsal views; anterior is up. Corresponding virtual transverse sections (along the yellow dotted lines) are shown in L’ and M’ respectively. Dorsal is up. Most EdU+ cells are *foxA*+ (L) or are encompassed by *calponin*+ smooth muscle tissues (M). In all relevant panels, white dashed lines correspond to the gut lining, while solid white lines delineate the outlines of the samples and white dotted lines correspond to the amputation planes. SX= segment number X.

To better describe the gut organization, we performed fluorescent immunostaining of acetylated tubulin, which labels the nervous system, cilia and musculature (Steinmetz et al., 2011; Planques et al., 2019), phalloidin F-actin staining which labels the musculature (Steinmetz et al., 2011) and microvilli (Tilic et al., 2023); and Hoechst DNA staining on histological sections (Supp. Fig. 1). It showed that the gut is composed of epithelial cells bearing microvilli, surrounded by an enteric nerve-net (Supp. Fig. 1A), the whole being encompassed by a sleeve of visceral smooth muscles (Supp. Fig. 1B). Thus, EdU+ cells labelled with a 5h-long pulse in S6/S7 are indeed located mostly inside the gut (Fig. 1C, C’), while in contrast, in S1 they were located not only in the gut but also in ectoderm and mesoderm (Fig. 1C’’). Meanwhile, cells labelled with a 48h-long EdU pulse are found in all tissue subtypes in S1, S6 and S7 (Fig. 1D to D”). We thus evidenced the presence of a gradient of cell proliferation along the anteroposterior (AP) axis of the uninjured worms in which proliferation rate decreases anteriorly but not in a homogenous way. Indeed, cell labelled with a rather short EdU pulse, referred as “quickly-cycling” cells, are found in all tissue subtypes in the posterior-most segment (S1) while they appear mostly restricted in the gut in more anterior segments (S6 and S7). The other tissues in anterior segments contain proliferative cells cycling more slowly.

Next, to determine the respective contribution of those quickly *versus* slowly-cycling cells to posterior regeneration, we incubated worms with EdU for 5h or 48h, as previously, and amputated them at two different positions (Fig. 1F). Either only the pygidium, growth zone and unrecognizable developing segments were removed (amputation “0”: the cut was done just behind S1), or five recognizable segments were removed in addition to the terminal part (amputation “−5”: the cut was made just behind S6). After 5 days of regeneration, the patterns of EdU+ cells in the regenerative parts were determined (Fig. 1G-J’). After a 5h-long EdU pulse followed by an amputation “−5”, only 9 % of the blastema cells were EdU+ (Fig. 1K) and appeared to be located mostly within the regenerating gut (Fig. 1G, G’). After a 5h-long EdU pulse followed by an amputation “0”, 19% of the blastema cells were EdU+ (Fig. 1K) and they were located not only in the gut but also in mesodermal and ectodermal tissues (Fig. 1H, H’). In contrast, after a 48h-long EdU pulse and whatever the type of amputation performed (amputation “−5” or “0”), more than 70% of the blastema cells were EdU+ (Fig. 1K) and located in all tissue types (Fig. 1I, I’ and J, J’). To confirm the particular contribution of quickly-cycling cells in anterior segments, to the regenerating gut, we coupled EdU labelling (*i.e.* after a 5h-long EdU pulse followed by an amputation “−5”) with *in situ* hybridizations for the gut specification factor *foxA* (Fig. 1L) and the smooth muscle marker *calponin* (Fig. 1M), (Brunet et al., 2016). The huge majority of the EdU+ cells colocalize with *foxA* inside the regenerating gut (Fig. 1L, L’) while only very scarce EdU+ cells colocalize with *calponin*, most of them being located inside the gut embedded in smooth muscles (Fig. 1M, M’).

Therefore, we described a gradient of proliferation along the AP axis of non-amputated *Platynereis* worms. Among those proliferative cells, we identified two types of progenitors (quickly-cycling *versus* slowly-cycling) which participate in posterior regeneration. Interestingly, almost all the blastema cells originate from slowly-cycling progenitors (labelled with a 48h-long pulse) in both anterior (S6/S7) and posterior (S1) segments. Moreover, quickly-cycling progenitors (labelled with a 5h-long EdU pulse) participate in the regeneration as well but to a lesser extent. In the posterior-most segment (S1), they produce derivatives in all cell compartments while in anterior segments (S6/S7), they participate mostly in the regeneration of gut cells. Those differences in patterns of EdU+ cells within the blastema reflects the initial patterns of quickly-cycling progenitors in the segments abutting the amputation plane, *i.e.* distributed in all cell compartments in the posterior-most segment and mostly localized within the gut in anterior segments.

### Tracing anterior gut cells with fluorescent beads demonstrates their lineage restriction through regeneration

To pursue our understanding regarding the contribution of those anterior quickly-cycling gut progenitors to the regeneration, we developed a means to specifically label all gut epithelial cells. We took advantage of their absorption property and tagged them with 1µm-diameter fluorescent beads. Indeed, after ingestion, the beads are specifically incorporated within the gut epithelial cells (Fig. 2A-A’’) to the exception of few posterior segments that were never labelled (Fig. 2A). With this new tool in hand, we then determined the participation of the gut epithelial cells in the regeneration. We amputated anteriorly worms which were labelled with fluorescent beads beforehand, and collected regenerative parts during the course of regeneration (5 stages) (Fig. 2B1-B5’). At stage 1 of regeneration when the wound epithelium has formed, we observed beads-labelled cells within the segment abutting the amputation where the gut has retracted (Fig. 2B1, B1’). As soon as the blastema has formed at stage 2 of regeneration (Fig. 2B2, B2’), we detected the fluorescent beads inside the regenerating gut consistent with the anus reformation. During the later growth and differentiation phases of regeneration (stages 3 to 5) the fluorescent beads’ signal remains restricted to the gut epithelium (Fig. 2B3-B5, 2B3’-B5’). At stage 5, we also combined the previous EdU labelling experiment (EdU+ cells in the regenerative part coming from the “quickly-cycling” gut progenitors, cf. Fig. 1G), with the use of fluorescent beads (Fig. 2C, C’). The overall colocalization of EdU+ cells and fluorescent beads demonstrates the congruence between the two technical approaches we have developed to label these gut cells as well as to determine their particular contribution to blastema gut cells.

**Figure 2:**
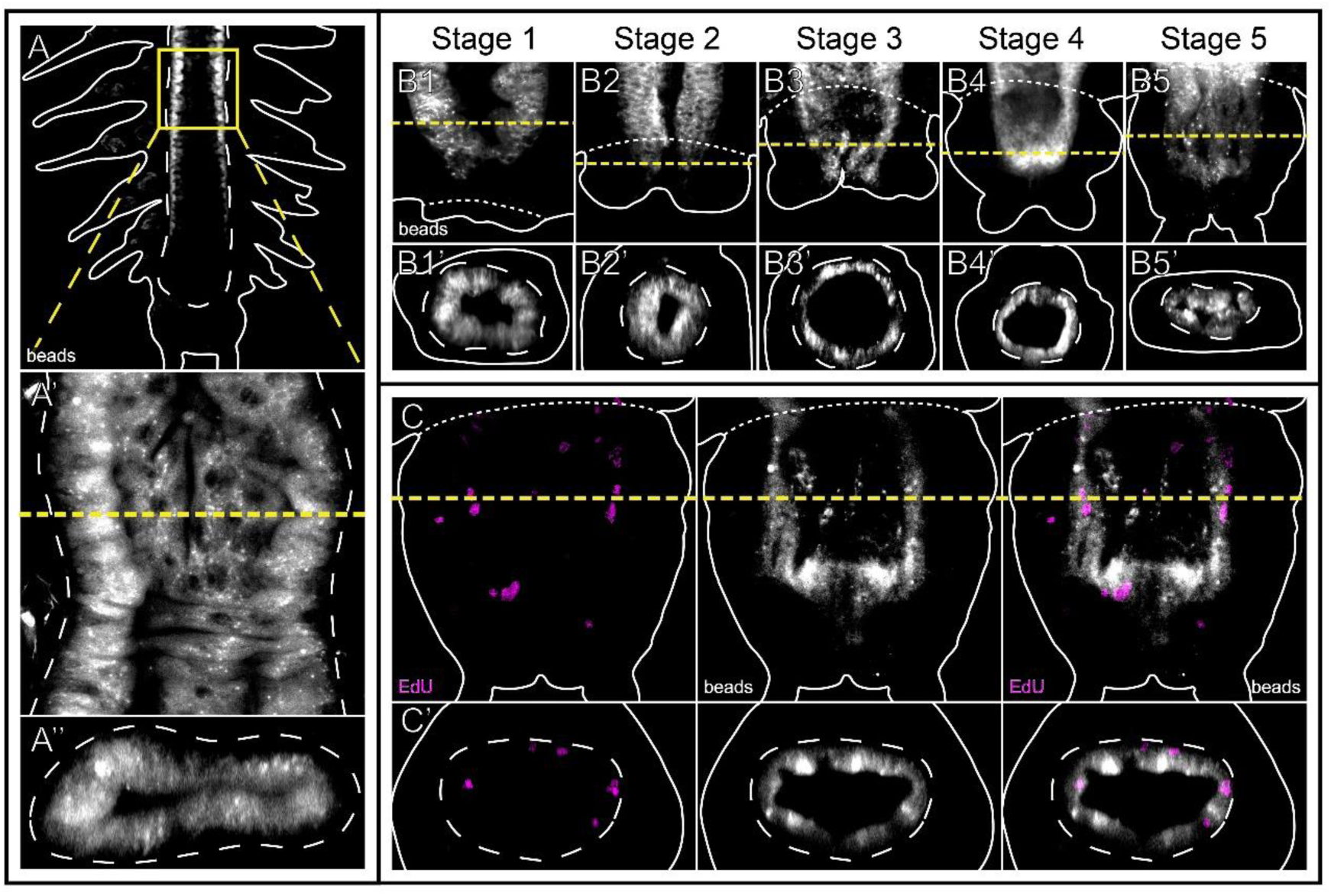
Gut progenitors localized anteriorly in the worm body are participating in regeneration and are lineage-restricted. **(A-C)** Unamputated juvenile worms were incubated with fluorescent beads for a week just before an anterior amputation and after a regeneration time of 1 to 5 days. **(A)** Confocal z-stack of the posterior part of a worm labelled with fluorescent beads (white) before amputation (dorsal view; anterior is up). A slight artefactual staining corresponding to parapodial glands is visible in some parapodia. **(A’)** Magnification focused on the gut along the yellow box in A. **(A’’)** Virtual transverse section along the yellow dotted line shown in A’ (Dorsal is up). The beads are located only in the gut epithelial cells. **(B1-B5)** Confocal z-stacks of worms labelled with fluorescent beads (white) at the indicated stages of regeneration. Dorsal views are shown on the top. Anterior is up. **(B1’-B5’)** Virtual transverse section along the yellow dotted lines are shown at the bottom. Dorsal is up. The fluorescent beads-labelled cells do participate in the regeneration of the gut. **(C)** Confocal z-stacks of a regenerative part at stage 5 after a dual fluorescent beads/EdU labelling (dorsal view, anterior is up). **(C’)** Virtual transverse section along the yellow dotted line shown in C. Dorsal is up. The cells proliferating before amputation participate mostly in the regeneration of the gut. In all relevant panels, white dashed lines correspond to the gut lining, while solid white lines delineate the outlines of the samples, yellow dotted lines indicate the virtual sections planes and white dotted lines correspond to the amputation planes.

Altogether, we conclude that gut progenitors which constitute a population of “quickly-cycling” cells inside anterior segments of unamputated worms, participate exclusively in the posterior regeneration of the gut epithelium and are thus lineage-restricted.

### Posterior gut cells are plastic and give rise to several cell lineages during regeneration

Next, we wondered whether the lineage-restriction of the anterior gut progenitors during regeneration also applies to the posterior-most segment. However, we had to overcome the fact that, during continuous growth, the most posterior gut epithelial cells are unable to incorporate fluorescent beads (Fig. 2A) and that a 5h-long EdU pulse labels many quickly-cycling progenitors outside the gut in posterior segments (Fig. 1A’, C’). A way to get over this is to take advantage of the fact that after an amputation in an anterior segment, “quickly-cycling” gut progenitors, labelled either with a 5h-long EdU pulse or with fluorescent beads, participate in the regeneration of gut cells which retain EdU and bead labellings when regeneration has finished (Fig. 1G, 2B5, 2C). Such regeneration procedures establish a proxy of posteriorized gut epithelial cells, which are labelled and can be tracked. Thus, we incubated unamputated worms either with fluorescent beads for a week, or with EdU for 5h, and amputated them anteriorly shortly after (Fig. 3A). We let the worms regenerate for 5 days (1^st^ regeneration, Cf. Fig. 1G) and performed a second amputation in the middle of the regenerated structure, removing the pygidium and growth zone that had regenerated after the first amputation. We let the worms regenerate a second time for a further 5 more days, performed a 1h-long BrdU pulse (only in the case of EdU incorporation) and collected the samples (2^nd^ regeneration, Fig. 3A).

**Figure 3:**
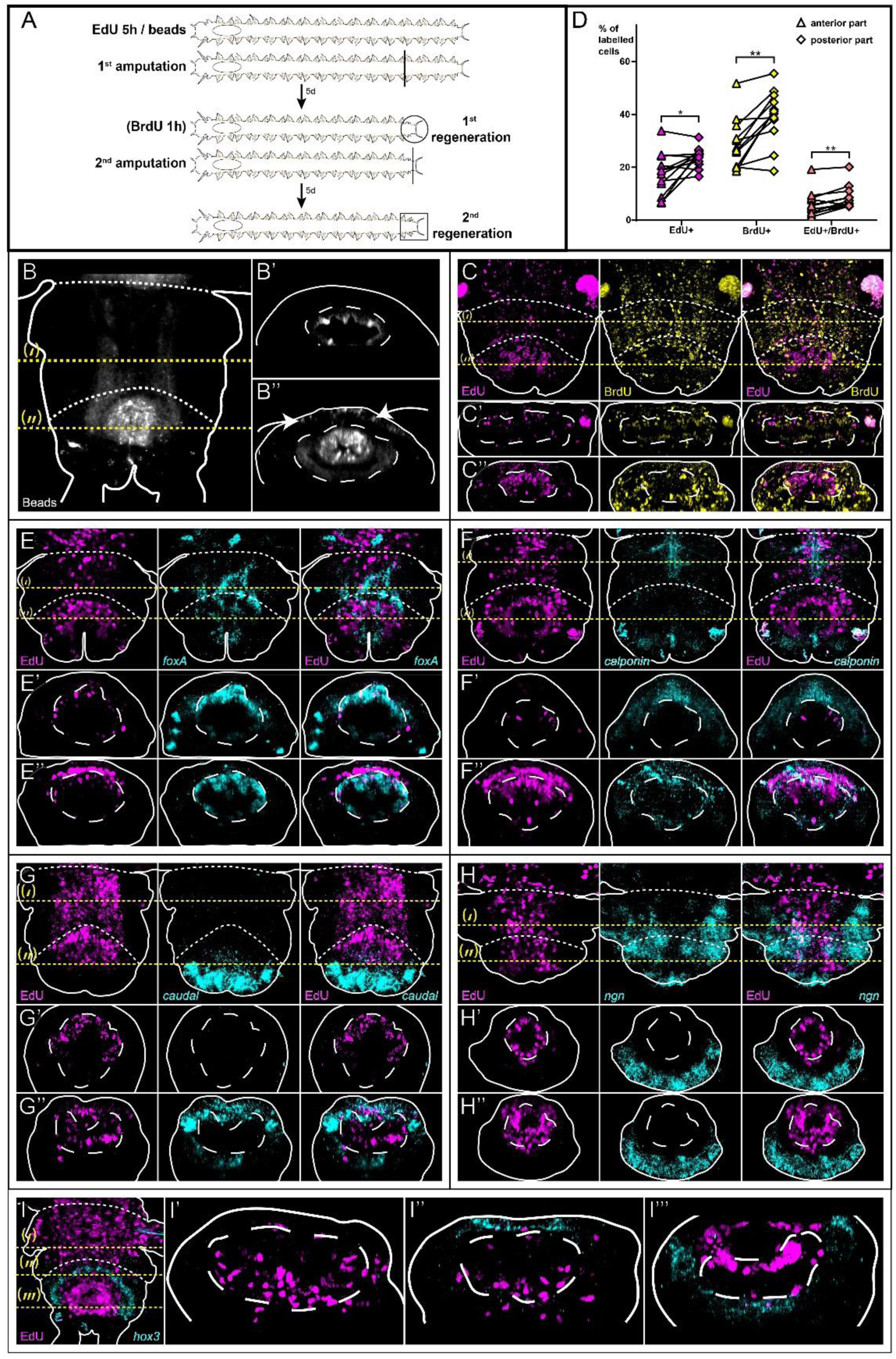
Gut progenitors localized posteriorly in the worm body are more plastic. **(A)** Schematic representation of the experiments performed: unamputated juvenile worms were incubated with fluorescent beads (for a week) or with EdU (for 5h) just before an anterior amputation. The worms were let to regenerate for 5 days and some of them were also incubated with BrdU (for 1h). The regenerative parts (1^st^ regeneration event, see Fig. 2) were re-amputated in the middle (*i.e.* removal of the regenerated pygidium and growth zone) and let to regenerate for 5 more days (2^nd^ regeneration event). **(B)** Confocal z-stack of a regenerative part obtained after both the amputations described in A were performed on worms labelled with fluorescent beads (white) (dorsal view; anterior is up). **(B’, B’’)** Virtual transverse sections along the yellow dotted lines (‘) and (‘’) in B (dorsal is up). **(B’)** In the anterior part of the samples (*i.e.*, remaining tissues of the 1^st^ regeneration), fluorescent beads are restricted inside the gut. **(B’’)** In the posterior part of the samples (tissues produced during the 2^nd^ regeneration), fluorescent beads are located both inside and outside the gut (white arrows). **(C)** Confocal z-stacks of a regenerative part obtained after both the amputations described in A were performed on worms incubated with EdU (magenta) and BrdU (yellow) (dorsal view; anterior is up). **(C’, C’’)** Virtual transverse sections along the yellow dotted lines (‘) and (‘’) shown in C (dorsal is up). **(C’)** In the anterior part of the samples (*i.e.*, remaining tissues of the 1^st^ regeneration) EdU+ cells are mostly located inside the gut. **(C”)** In the posterior part of the samples (tissues produced during the 2^nd^ regeneration), EdU+ cells are located inside and outside the gut. **(D)** Comparison of the proportions of EdU+, BrdU+ and EdU+/BrdU+ cells between the anterior parts (triangle) and the posterior parts of the samples (diamond) from C (Wilcoxon signed-rank test: * p <0.05; ** p <0.01). **(E-I)** Confocal z-stacks of regenerative parts obtained after both the amputations described in A were performed on worms incubated with EdU (dorsal views, anterior is up). *In situ* hybridization signals (cyan) for the gut (*foxA,* (E)), the smooth muscle (*calponin,* (F)), the pygidium (*caudal,* (G)), the neural (*neurogenin or ngn,* (H)), or the growth zone (*hox3*, (I)) markers. **(E’-H’, E’’-H’’)** Virtual transverse sections along the yellow dotted lines (‘) and (‘’) shown in E – H (dorsal is up). In the anterior part of the samples (E’), most EdU+ cells are located inside the gut (and colocalize with *foxA*) while a significant amount of them are outside the gut in the posterior part (E’’). Those EdU+ cells colocalize with *calponin* and *caudal* (F’’, G’’) but not ventrally with *ngn* (H’’). **(I’-I’’’)** Virtual transverse sections along the yellow dotted lines (‘), (‘’), (‘’’) shown in I (dorsal is up). Very few EdU+ cells colocalize with *hox3* dorsally (I’’) and ventrally (I’’’) in the regenerated growth zone. In all relevant panels, white dashed lines correspond to the gut lining, while solid white lines delineate the outlines of the samples, yellow dotted lines indicate the virtual sections plans and white dotted lines correspond to the amputation planes.

We observed differences in the distribution of fluorescent beads-labelled (Fig. 3B) or EdU+ cells (Fig. 3C) along the anteroposterior axis of the “blastema-like” structure (*i.e.* a structure composed of a mix of a 10 dpa (anteriorly) and a 5 dpa (posteriorly) regenerated structure). In the anterior part, which corresponds to the remaining tissues of the 1^st^ regeneration, the fluorescent beads and the EdU+ cells were still specifically or mainly restricted to the gut, respectively (Fig. 3B’, C’). Strikingly, in the posterior part of the blastema-like structure, which corresponds specifically to the 2^nd^ regeneration event, fluorescent beads and EdU+ cells were detected both inside but also outside the gut, mainly in the dorsal side of the regenerating part (Fig. 3B’’, C’’). Furthermore, there were significantly more EdU+ cells in the posterior part (23.8%, Fig. 3D) than in the anterior part (17.4%, Fig. 3D). Similarly, the number of BrdU+ and EdU+/BrdU+ cells were higher in the posterior part of the structure than in the anterior part (Fig. 3D). This means that the gut progenitors, which were previously labelled with EdU, are actively mobilized during the 2^nd^ regeneration to participate to the formation of seemingly other tissues than the gut, and more than a third of them are still proliferative. In addition, highly-dividing cells participating in the 1^st^ regeneration labelled with BrdU, contribute as well massively to this 2^nd^ event of regeneration (Fig. 3C to C’’, D)

Then, we determined the molecular identity of the cells originating from the gut progenitors that we observed outside the gut after the 2^nd^ regeneration (Fig. 3B’’, C’’) by coupling EdU labelling and *in situ* hybridizations for tissue-specific genes (Fig. 3E-I’’’). It included the gut specification factor *foxA* (Fig. 3E to E”), the smooth muscle marker *calponin* (Fig. 3F to F”), the pygidial marker *caudal* (Fig. 3G to G”), the neural progenitor factor *neurogenin* (Fig. 3H to H”) and the ectodermal growth zone stem cells marker *hox3* (Fig. 3I to I”’). In the anterior part of the “blastema-like” structure, EdU+ cells mostly colocalize with *foxA* endodermal expression and are encompassed by the mesodermal expression domain of *calponin*, similarly to the patterns obtained earlier for the 1^st^ regeneration (Fig. 3E’, F’; Cf. Fig. 1L-M’). In contrast, in the posterior part of the “blastema-like” structure, many EdU+ cells did not colocalize with *foxA* but did with *calponin* (Fig. 3E’’, F’’). Some of the EdU+ cells expressed also *caudal* within the mesoderm and the ectoderm of the regenerating pygidium (Fig. 3G’’). In contrast, we never observed any colocalization with the *neurogenin* expression domain (Fig. 3H’’) in the ventral neurectoderm. Similarly, the EdU+ cells did not colocalize with the ventral part of *hox3* expression domain in the ectodermal growth zone (Fig. 3I, I’’’). In the dorsal part of *hox3* expression domain, the presence of very few EdU+ cells cannot be ruled out (Fig. 3I’’).

Taken together, those results showed that the quickly-cycling gut progenitors, which were only able to regenerate gut during a previous anterior amputation, can give rise to other tissues after posteriorization. While they can be directed towards an epidermal, mesodermal and posterior ectodermal fate, they cannot give rise to neural progenitors and probably not to stem cells from the ectodermal growth zone. Those gut cells have therefore acquired plasticity upon posteriorization. These results support the idea that the gut progenitors within the most posterior segments could display such plasticity compared to their anterior lineage-restricted counter parts. It suggests the presence along the AP axis of a gradient of cellular plasticity positively associated with the gradient of proliferation of the tissues we evidenced previously.

### Cell migration and proliferation as well as tissue maturity regulate gut cell plasticity

This intriguing plasticity harboured by posterior gut progenitors is *de facto* spatially limited. Indeed, we showed that anteriorly-located gut progenitors are lineage-restricted. Their plasticity is thus lost upon growth and differentiation of the tissues. We aimed here at describing the cellular mechanisms underlying this plasticity; and also, at determining at which point the posteriorized gut progenitors would lose it. As cell proliferation and migration are often required in similar processes (Friedl and Gilmour, 2009), we hypothesized it would be the case here as well. To this end, we performed the same reamputation procedure, but this time, we incubated the worms with cell proliferation (Hydroxy-Urea or HU) or migration, through actin polymerisation, (LatrunculinB or LatB) widely used inhibitors during the second phase of regeneration (Fig. 4A). Previous results showed that the inhibition of cell proliferation with HU does not prevent the formation of the blastema but hinders its growth and differentiation (Planques et al., 2019). The inhibition of cell migration (and modifications of cell shape) with LatB slows down regeneration and causes mild morphological defects (thicker anal cirri, Supp. Fig. 2). In contrast to the controls in which EdU+ cells are found, as expected, outside the gut in the posterior region of the blastema-like structure (Fig. 4B-B”), when inhibiting cell migration with LatB, most EdU+ cells stayed mostly restricted inside the gut in both anterior and posterior parts of the “blastema-like” structure (Fig. 4C to C’’). Similarly, most EdU+ cells are restricted inside the gut in the “blastema-like” structure when inhibiting cell proliferation with HU (Fig. 4D-D’’). Next, to determine more precisely at which point the posteriorized gut progenitors lose their enhanced plasticity, we performed the reamputation procedure, but at later stages of initial regeneration (*i.e.* at 9dpa or 12dpa, instead of 5dpa) when newly produced tissues are differentiating (Fig. 4A). We observed that, as expected, the distribution of EdU+ cells is mostly restricted inside the gut in the anterior part of the regenerating structure (1^st^ regeneration, Fig. 4 E, E’, F, F’) but also in its posterior part (2^nd^ regeneration, Fig. 4E, E’’, F, F’’), showing that posteriorized gut progenitors do no longer harbour plasticity at those stages.

**Figure 4:**
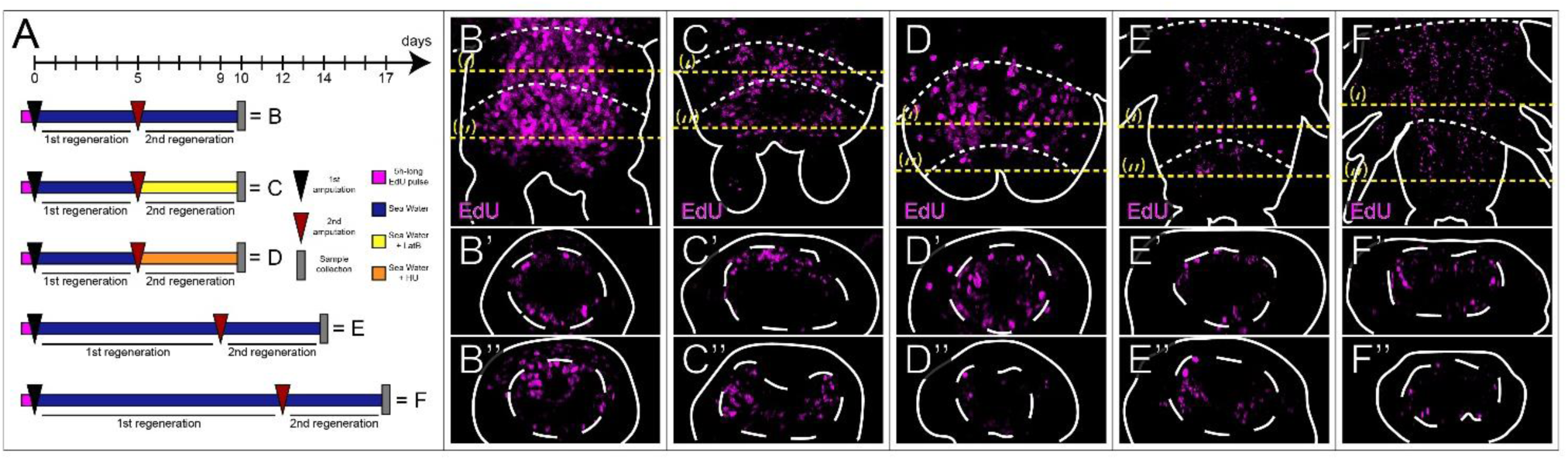
Cell proliferation and migration, as well as tissue maturity modulate the plasticity of posteriorized gut progenitors through regeneration. **(A)** Schematic representation of the experiments performed: all worms were incubated in EdU for 5h prior to a 1^st^ amputation “−5“ (5 segments and the pygidium were removed). After 5, 9 or 12 dpa, a 2^nd^ amputation was performed in the middle of the regenerated structure. The second regeneration event was either performed in sea water (B, control) or in the presence of pharmacological inhibitors (C, LatrunculinB or LatB; D, Hydroxy-Urea or HU). **(B-F)** Confocal z-stacks of regenerative parts obtained after both the amputations and the different incubations described in A are shown on the top (dorsal views, anterior is up). **(B’-F’)** and **(B’’-F’’)** are virtual transverse sections along the yellow dotted lines (‘) and (‘’) shown in B-E (dorsal is up). In the posterior part of the samples (C’’ to F’’), EdU+ cells (magenta) are mostly located inside the gut, while they are also present outside the gut in the control (B’’). In all relevant panels, white dashed lines correspond to the gut lining, while solid white lines delineate the outlines of the samples, yellow dotted lines indicate the virtual sections plans and white dotted lines correspond to the amputation planes.

Hence, the reorganisation of the actin filaments and its related cellular functions (changes in cell shape and migration), as well as cell proliferation are crucial for the posteriorized gut progenitors to acquire their plasticity. Besides, this plasticity is lost very quickly upon maturation of the tissues.

### Most of the blastema cells are from local origin, including the regenerated posterior stem cells

In the previous sections, we have shown that, whatever their position along the AP axis of the worms, the gut progenitors only partly contribute to the whole blastema formation, even the plastic posteriorized gut progenitors. Indeed, the anterior gut progenitors provide blastema gut cells, and the posterior ones, while producing several derivatives cannot regenerate neural tissues (Fig. 3H) and can only at best regenerate few cells of the ectodermal growth zone stem cells (Fig. 3I). This led us to wonder about the origin of the cells that will give rise to the rest of the tissues, in particular, the stem cells of the regenerated growth zone. As the position along the AP axis of the gut progenitors impacts their fate, we also ask ourselves if this could be the case for the other cells participating in the blastema formation. Our previous work proved that most of the blastema cells are from local origin, from the segment abutting the amputation plane, in a context of an anterior amputation (*i.e.* amputation “−5”, (Planques et al., 2019)). We thus sought to determine whether the source of the regenerative cells would remain local along the anteroposterior axis and also whether the stem cells of the growth zone would be from local origin as well. To discriminate the different involvement of local *versus* more distant tissues to the regeneration along the body axis, we used an experimental set-up consisting in two serial amputations performed either in anterior (“−5”) (as in (Planques et al., 2019)) or posterior (“0”) locations (Fig. 5 and Supp. Fig. 4; and Supp. Fig. 3, respectively). First, to have a proxy of the cells activated by anterior or posterior amputation, we performed a 5h-long EdU pulse at stage 1 when regeneration has been initiated and the wound epithelium formed (Fig. 5A), chased the EdU for two days (*i.e.* until 3 dpa) and performed a 1h-long BrdU pulse before collecting the samples. We determined that 52.8% of the blastema cells (at 3dpa for an anterior amputation) arose from those EdU+ cells proliferating after amputation (Fig. 5B). Besides, they contribute massively to the pool of highly proliferative cells at 3dpa (among the 31.8 % of BrdU+ cells, 80% of them are also EdU+), which means that those EdU+ cells participate not only in the formation of the blastema but also in the following phases of regeneration. Thus, they harbour a high regenerative potential. We aimed at quantifying the precise difference of regenerative potential of the EdU+ cells in the segment abutting the amputation plan with those in the segment directly anterior to it, for the two positions of amputations (anterior *versus* posterior). To this extent, as previously, we amputated worms anteriorly (see Fig. 5) or posteriorly (see Supp. Fig. 3) and let them regenerate for 24h, performed a 5h-long EdU pulse to label the cells activated by the amputation and let the worms regenerate two more days when the blastema has formed. Then, we either amputated the blastema and the first abutting segment (Amputation A) or only the blastema (Amputation B). Following this second amputation, we let the worms regenerate three more days until another blastema has formed, and performed a 1h-long BrdU pulse before collecting the samples to quantify how many EdU+ are still proliferative after both amputations (Fig. 5A).

**Figure 5:**
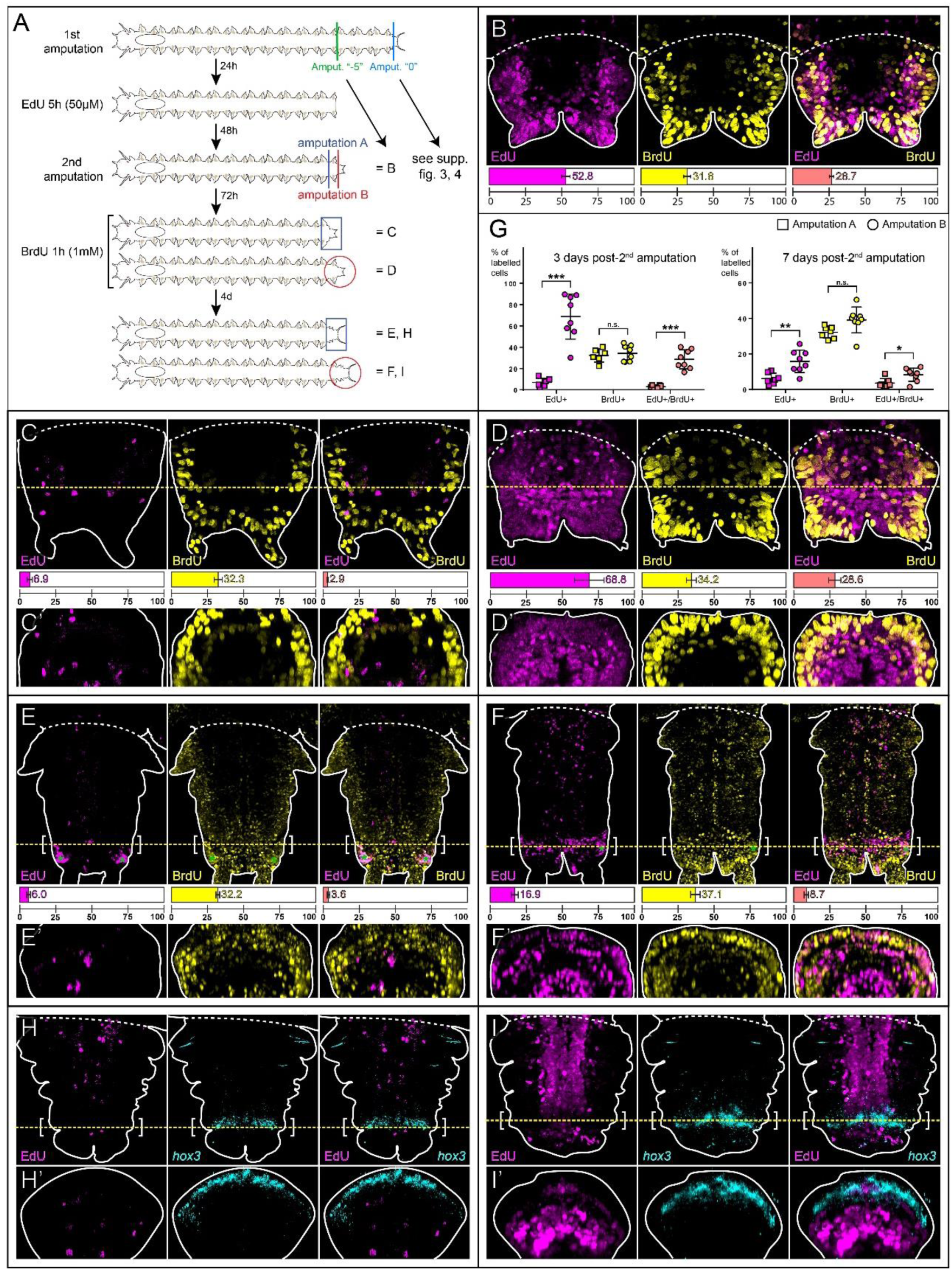
Most of the blastema cells originate from the segment abutting the amputation including the stem cells of the regenerated growth zone. **(A)** Schematic representation of the experiments performed: all worms were amputated anteriorly (amputation “−5”: 5 segments and the pygidium were removed) and let to regenerate for 24h, then incubated with EdU for 5h and let to regenerate for an additional 48h. Some of them were incorporated with BrdU (for 1h) (B). The other worms were then re-amputated: either the blastema and the abutting segment were removed (=amputation A) (C, E and H), or only the blastema was removed (=amputation B) (D, F and I). The worms were then let to regenerate for three days, incubated with BrdU (for 1h) and collected (C and D) or let to regenerate for four more days (E and F). Others were just let to regenerate for seven days after the 2^nd^ amputation (H and I). **(B)** Confocal z-stacks of a regenerative part obtained after an amputation “−5”, incubated with EdU (magenta) at 1dpa and BrdU (yellow) at 3dpa (ventral view, anterior is up). **(C-F)** Confocal z-stacks of regenerative parts obtained after both the amputations described in A were performed on worms incubated with EdU (magenta) and BrdU (yellow), (ventral views; anterior is up). **(C’-F’)** Virtual transverse sections along the yellow dotted lines shown in C-F (ventral is up). **(G)** Comparisons of the proportions of EdU+, BrdU+ and EdU+/BrdU+ cells between samples at 3 days post-2^nd^ amputation (C and D) on the left and at 7 days post-2^nd^ amputation (E and F) on the right (square = amputation A; circle = amputation B). Each bracket corresponds to a Mann-Whitney U test. n.s. p >0.05; * p <0.05; ** p <0.01; *** p<0.001. Mean + SD are shown. EdU+ and EdU+/BrdU+ cell proportions are significantly higher in D than in C, and in F than in E. **(H-I)** Confocal z-stacks of regenerative parts at 7 days post-2^nd^ amputation showing EdU+ cells (in magenta) as well as *in situ* hybridization signal (in cyan) for *hox3* (ventral views, anterior is up). Corresponding virtual transverse sections (along the yellow dotted lines) are shown in H’ and I’ (ventral is up). EdU+ cells colocalize with *hox3* in I but do not in H. In all relevant panels, solid white lines delineate the outlines of the samples, yellow dotted lines indicate the virtual sections plans and white dotted lines correspond to the amputation planes. White brackets indicate the growth zone and green asterisks indicate artefactual staining corresponding to pygidial glands.

First, we observed that the number and localization of BrdU+ cells are similar between condition A (Fig. 5C, Supp. Fig 3A) and B (Fig. 5D, Supp. Fig 3B; roughly 33 %, see individual values in Fig. 5G and Supp. Fig 3C), whatever the position of the amputation (anterior *versus* posterior). It is consistent with the fact that the BrdU pulse has been performed at the same stage of regeneration (*i.e.* 3 days after the second amputation), demonstrating that the different serial amputations do not have significant effects on proliferation profile. In contrast, we obtained very distinct EdU+ cell patterns in the blastema depending on the second amputation condition (A *versus* B). For an anterior amputation, in condition A, only around 7% of internal cells are EdU+ (Fig. 5C, C’, G) whereas around 69 % of the blastema cells are EdU+, in both superficial and internal tissues, in condition B (Fig. 5D, D’, G). The situation is rather similar for a posterior amputation: while in condition A, around 27 % are EdU+ (Supp. Fig. 3A, A’, E), around 75 % of the blastema cells are EdU+ in condition B (Supp. Fig. 2B, B’, E). Finally, looking at the EdU+/BrdU+ cells, there is obviously a huge difference in proportions between conditions A and B accounting for the fact that there are 3 to 10-fold more EdU+ cells in B than in A (for both posterior and anterior amputations respectively). However, interestingly, roughly 40% of the EdU+ cells are BrdU+ at 3 days post-2^nd^ amputation in all conditions (Fig. 5C, D, G; Supp. Fig. 3A, B, E).

Given that the only difference between conditions A and B is the absence, or the presence, of the segment abutting the first amputation plane, we conclude from these experiments that the cells activated by the amputation and located close to the wound have a higher plasticity than the ones located more anteriorly. Thus, they contribute to most of the blastema cells, regardless the site of the initial amputation (anterior *versus* posterior).

Next, we sought to determine the origin of the stem cells of the regenerated growth zone, and more precisely whether they would be from local origin as well, all along the AP axis. To track the regenerated stem cells, we relied on the fact that they constitute a population of Label Retaining Cells (LRC, (de Rosa et al., 2005)). Indeed, as putative stem cells, their proliferation rate is rather low and consequently after an EdU incorporation the signal should be retained longer than in other cell types. To this end, we used the same experimental set-up as described previously, but instead of collecting the samples after the 1h-long BrdU pulse, we chased both the EdU and BrdU by letting the worms regenerate for four more days, until the very end of regeneration process (Fig. 5A). For an anterior amputation, as expected, the pattern of BrdU+ cells is similar between conditions A and B (Fig. 5E, E’, F, F’ and G). In contrast, the patterns of EdU+ cells are very distinct between both conditions. In condition A, there are still very few EdU+ cells (6%, Fig. 5G) scattered in internal tissues (Fig. 5E, E’). As for condition B, there is a dramatic 4-fold reduction of the proportion of EdU+ cells in the samples (from around 69 % to 17 %, Fig. 5F, G). This reflects the fact that most of the blastema cells underwent enough cell divisions for the EdU to be diluted. Those remaining 17 % of EdU+ cells appear localized for few of them inside the gut but more importantly at the interface between the pygidium (the terminal part of the worm body) and the growing new segments which presumably corresponds to the regenerated growth zone stem cells (Fig. 5F, F’). This EdU pattern is reminiscent of what could be expected for an LRC. The similar experimental approach has been followed after an initial posterior amputation (Fig. 5A). In this case, there are higher proportions of EdU+ cells in both conditions (17.9% for condition A and 36.2% for condition B, Supp. Fig. 3C, D, E) but similarly EdU+ cells do not appear to be located in the growth zone in condition A whereas they do in condition B (Supp. Fig. 3C-D’). The higher proportions of EdU+ cells for both conditions (A and B) for an amputation “0”, in comparison to an amputation “−5” may reflect that the tissues abutting a first initial posterior amputation are more proliferating than more anterior tissues. To confirm the localization and identity of those LRC, we coupled EdU labelling with whole-mount *in situ* hybridizations for markers of different cell populations of the growth zone (Gazave et al., 2013). We selected the genes *hox3* and *evx* – expressed in the ectodermal cells of the growth zone, as well as *piwiB*, expressed in both the ectodermal and mesodermal cells of the growth zone (Fig. 5 and Supp. Fig. 4). After an initial anterior amputation, in condition A, the few remaining internal EdU+ cells do not colocalize with *hox3* signal (Fig. 5H, H’) whereas they do colocalize in condition B (Fig. 5I, I’). Similar results are obtained for *evx* (Supp. Fig. 4A-B’) and *piwi* (Supp. Fig. 4C-D’).

Those results demonstrate that the stem cells of the regenerated growth zone originate from local cells (*i.e.* from the segment abutting the amputation plane) activated by the amputation, whether this amputation has been performed anteriorly or posteriorly along the animal body axis. In addition, those cells have a higher plasticity compared to those located more distantly to the amputation plan (*i.e* one segment upstream).

## Discussion

### Homeostatic progenitors with accelerated cell cycle are a cellular fuel for regeneration

Our proliferation labelling experiments revealed the contribution of two types of progenitors (quickly-cycling *versus* slowly-cycling) from non-amputated tissues in *Platynereis* posterior regeneration. Importantly, the slowly-cycling progenitors give rise to the huge majority of the blastema cells. Slowly-cycling stem/progenitor cells are the main source of regenerative cells in many other contexts of regeneration for a diversity of species. In mammalian skin (Koren et al., 2022) and gut (Karmakar et al., 2020) regeneration, they are critical for wound repair while their role during homeostasis may differ. Similarly, slowly-cycling stem/progenitor cells are also involved in higher scale of regeneration, such as whole-body regeneration in planarians (Molinaro et al., 2021) or in the ctenophore *Mnemiopsis leidyi* (Ramon-Mateu et al., 2019), as well as in head regeneration in the cnidarian *Hydra* (Govindasamy et al., 2014). Usually, the cell cycle of those slowly-cycling progenitors has to accelerate to produce the cells required for regeneration. This can be achieved through, either, the acceleration of their S phase (such as in *Drosophila* imaginal disc regeneration, (Crucianelli et al., 2022)) or the shortening of their G1 phase (such as in *Drosophila* gut regeneration, (Cohen et al., 2021) or spinal cord regeneration in axolotl (Cura Costa et al., 2021)).

In *Platynereis*, the replication rate of the slowly-cycling progenitors does not match the timing of posterior regeneration. Indeed, it requires 48 hours to label those progenitors with EdU whereas for the same duration, a small highly proliferative blastema has already formed (Planques et al., 2019). Thus, only a cell-cycle acceleration can explain their massive contribution to the blastema. In support to this assumption, a recent study on another annelid species reports the overall shortening of the cell cycle upon amputation (Shalaeva and Kozin, 2023). A precise determination of the cell cycle parameters during regeneration in *Platynereis* remains to be performed, but the identification of those two types of progenitors was decisive for a better understanding of the cellular sources of posterior regeneration in *Platynereis*.

### A cellular model for posterior regeneration in *Platynereis*

In this study, we aim to define a model of regeneration for the annelid *Platynereis dumerilii* at the cellular level. We first obtained new insights on the fate of the cells participating in the regeneration. We highlighted a gradient of plasticity of gut progenitors through regeneration by showing that, in contrast to their lineage-restricted anterior counterparts, posterior gut progenitors can widen their progeny during regeneration and produce ecto/mesodermal derivatives. This gradient of plasticity was shown to be positively associated with a gradient of proliferation of the tissues, with the most proliferative tissues being localized posteriorly. As those posterior-most tissues close to the growth zone are also the most-recently formed, we can easily presume that anterior tissues are overall more differentiated than posterior ones. This intuitive assumption for a continuously growing animal is supported by scattered data. For instance, posterior-most segments never harbour chaetae (extracellular structures produced by mature parapodia) nor express a chaetae-associated gene marker (*Chitin Synthase*) (Gazave et al., 2017). Similarly, they do not contribute to gas exchanges, as they never express globin markers (Song et al., 2020), nor to an efficient digestion, as highlighted by their inability to incorporate fluorescent bead (Fig. 2A). Thus, the gradients of plasticity and proliferation we evidenced can be negatively associated with a gradient of tissue differentiation and as such, we can link the plasticity of posterior gut progenitors to their low level of differentiation.

Moreover, we went further into the origin of the blastema cells. It was shown previously in *Platynereis* that most of the blastema cells are from local origin, arising from the segment directly abutting the amputation plane (Planques et al., 2019). We confirmed this finding by determining that most blastema cells, including the stem cells of the regenerated growth zone, are from local origin whatever the level of differentiation of the segment abutting the amputation plane (*i.e.* anterior or posterior tissues). Comparing the origin(s) of the cells participating in the regeneration of *Platynereis* with what is known in other annelid species (Bely, 2014) and beyond is mandatory to determine the diversity of cellular processes at play during regeneration in animals and understand its evolution (Grillo et al., 2016). Two main cellular models prevail for blastema formation in animals (Bideau et al., 2021). First, in species harbouring extensive regeneration ability, notably whole-body regeneration, such as the planarian *Schmidtea mediterrannea* or the cnidarian *Hydractinia symbiolongicarpus*, regeneration processes rely on migrating adult pluripotent stem cells able to produce both somatic and, for the latter, also germinal derivatives (Wagner et al., 2011; Varley et al., 2023). The presence of adult stem cells (pluripotent or with less potency) participating in annelid regeneration was often proposed, while never properly evidenced, in many annelids, such as earthworms (*Lumbriculus* sp. and *Enchytraeus japonensis*, (Randolph, 1892; Myohara et al., 1999; Sugio et al., 2012), or the capitellidae *Capitella teleta,* (Sedentaria, (de Jong and Seaver, 2017)). In addition, a recent single cell atlas for another sedentaria species, *Pristina leidyi* suggests the presence of such stem cells with a pluripotency signature, but their putative role during regeneration remains to be established (Álvarez-Campos et al., 2023). While we cannot definitely rule out this possibility, our data does not support the idea that adult multi/pluripotent stem cells are involved during posterior regeneration in *Platynereis*. The second main cellular model relies mostly on dedifferentiating lineage-restricted cells close to the wound (with sometimes also the involvement of tissue resident lineage-restricted stem cells), *e.g.* in the salamander limb regeneration (Kragl et al., 2009). Dedifferentiation was also proposed for a couple of annelid species (*i.e.* the errantia *Syllis malaquini* (Ribeiro et al., 2021) or *Alitta virens* (Shalaeva and Kozin, 2023)) during posterior regeneration. So far, the cellular trajectories supporting this idea remain elusive but such data already highlight the likely diversity of cellular mechanisms of regeneration within annelids (Bely, 2014).

In sum, our data led us to define a hybrid cellular model (summarized in Fig. 6) for *Platynereis* posterior regeneration: most of its regenerative cells are from local origin (like in salamanders) but the massive contribution of slowly-cycling progenitors rules out the possibility of a “complete” dedifferentiation process in which post-mitotic cells re-enter cell cycle. This rather supports the idea of pools of specialized progenitors with different replication rates, maintained throughout juvenile stage, that are mobilized upon amputation trigger. Among them, posterior gut progenitors can become plastic during regeneration.

**Figure 6:**
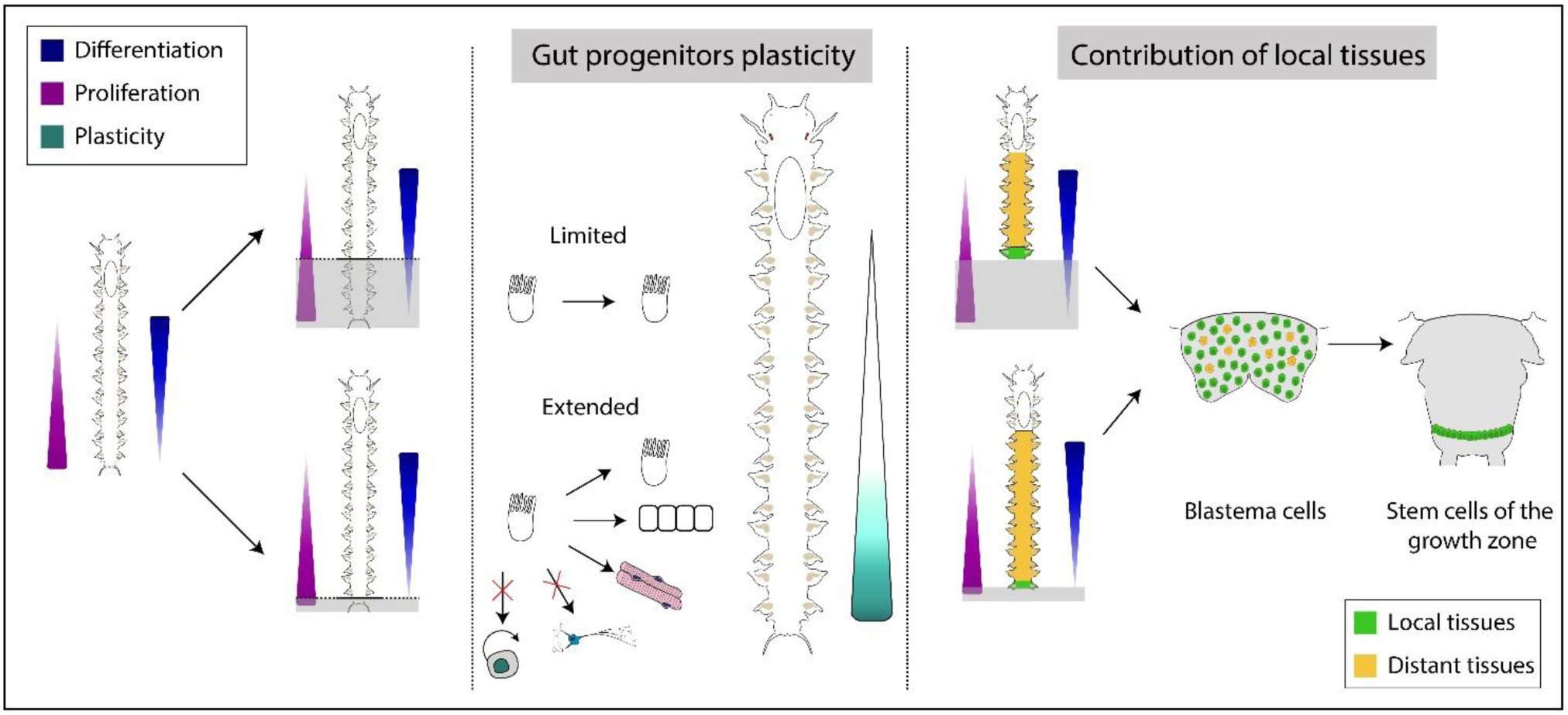
Cellular model for posterior regeneration in the annelid Platynereis dumerilii. Opposite gradients of cell proliferation and differentiation shape tissue maturity and gut plasticity along the antero-posterior (AP) axis of the worms. Upon anterior amputation, gut progenitors are lineage restricted and produce only gut epithelial cells. Upon posteriorization, gut progenitors contribute to ectodermal and mesodermal tissues in addition to gut epithelial cells, thus defining a gradient of plasticity. Gut progenitors’ plasticity is however limited as no posterior stem cells nor neural derivatives are produced upon metaplasia. Whatever the position of the amputation along the AP axis, the contribution of tissues abutting the amputation plan is massive for the blastema formation and the growth zone stem cells regeneration. Gradient of cell proliferation is depicted in magenta, while an opposite gradient of tissue differentiation is in blue. Gradient of gut progenitors’ plasticity is in blue-green. Local and distant tissues are depicted in green and yellow respectively.

### Plasticity of gut progenitors in *Platynereis* as an example of metaplasia during regeneration

Historical seminal studies of embryonic development led to the formalization of a broad model in which cell differentiation is an irreversible process, and development as a whole constitutes a gradual loss of potency from the totipotent zygote to fully differentiated adult cells (Caplan and Ordahl, 1978; Merrell and Stanger, 2016). This inflexibility of cell identity is nowadays reconsidered in the light of recent genetic lineage tracing experiments, which uncover various developmental processes showing metaplasia *i.e.* the acquisition of a cell identity that is unusual for a given tissue (Virchow, 1886; Mills et al., 2019). Metaplasia was often observed during various regeneration contexts in which progenitor/stem cells can acquire a higher plasticity compared to tissue homeostasis, even though it is always limited in terms of potency and temporality. For instance, in the zebrafish fin, specific subpopulations of fibroblasts can restore more types of fibroblasts during regeneration than they do during homeostasis, but they can only produce fibroblasts, showing that the expansion of their potency is very limited (Tornini et al., 2017). Likewise, in mammals, a skin injury can transiently widen the progeny of specific skin stem cells which can then produce all the diversity of skin cells but not cells outside the skin (Blanpain and Fuchs, 2014).

In this study, we uncovered the intriguing ability for posterior less differentiated gut progenitors to produce other types of derivatives (*e.g.* epidermis or muscle) during regeneration, which can be considered as an example of metaplasia as well. Interestingly, the metaplasia of the gut in *Platynereis* is also limited in different ways. It is rather transitory; metaplasia probably stops as soon as the gut starts to differentiate. Related to this, it is also limited spatially in the body of the animal: only newly-produced and thus posterior gut progenitors can switch lineage during regeneration. Besides, it is limited in potential: the extent of the metaplasia is not total as posterior gut progenitors cannot produce nervous system or putative posterior stem cells.

Why are posterior gut progenitors with enhanced plasticity, unable to produce nervous system derivatives upon amputation? In many annelids, the ventral nerve chord (VNC) from non-amputated tissues plays a major role on the reformation of the nervous system in the regenerated structure (Sinigaglia and Averof, 2019). In *Platynereis*, nerves from the VNC will rapidly innervate the blastema (Planques et al., 2019) and may serve as both direct or indirect source of signals (*e.g.* in the absence of the VNC, an aneurogenic regenerating structure is produced in a sister-species (Boilly et al., 2017)). We can speculate that such posterior gut progenitors cannot replace this important signal as well as the physical support provided by the VNC, in contrast to muscle cells which appear *de novo* in the blastema and do not seem to rely on the muscle system in non-amputated tissues for their initial differentiation (Planques et al., 2019). Upon amputation, posterior gut progenitors also never produce the putative posterior stem cells responsible for the constant growth of the worms (Gazave et al., 2013). We can imagine that “enhanced” posterior gut progenitors are unable to produce stem cells as the latter may harbour a too high potency, unreachable through their metaplasia. However, our results indicate that other cells located in the segment abutting the amputation plane and whose identity remains to be determined can reach such potency to produce posterior stem cells. As such, the gut is not the only tissue capable of metaplasia following the amputation signal. That is why it would be of special interest to follow the fate of other tissues (*e.g.* muscles or nervous system). The development of genetic lineage tracking would allow to determine whether this metaplasia is specific to given types of tissue or if it depends solely on the differentiation level of any tissue.

### Posterior regeneration in *Platynereis* is a plastic process at the cellular level but is morphologically robust

This phenomenon of metaplasia of posterior gut cells highlights an important point regarding the cellular processes involved during regeneration. Indeed, different sources of cells along the antero-posterior axis of the animal contribute to the blastema and eventually to a morphologically identical structure. This notion of different cellular “paths” for a same regeneration process is intriguing and rather uncommon. One key akin example is the retinal regeneration of *Xenopus* in which the involvement of three different cellular sources has been reported, depending on the extent of the injury (Parain et al., 2023). So far, we have no idea about the molecular mechanisms underlying such diversity of cellular mechanisms for regeneration. We recently unravelled the transcriptional landscape of posterior regeneration in *Platynereis* (Paré et al., 2023), following an “anterior” amputation and it would be interesting to obtain comparable data for a more posterior amputation. In addition, those results led us to consider the robustness of the posterior regeneration in *Platynereis*. In a previous study, we reported that serial amputations (up to 10) do not impair regeneration efficiency, at least at the morphological level (Planques et al., 2019). Similar robust regeneration events after successive injuries exist in some species, such as the zebrafish fin (Azevedo et al., 2011) and the newt’s lens that can properly regenerate even after 18 repeated lens removal spanning over 30 years (Eguchi et al., 2011), without any modification of the associated transcriptional program (Sousounis et al., 2015). In contrast, in many other species, the succession of injuries alters regeneration effectiveness (Eming et al., 2014) by either exhausting the tissue-specific stem cell pool (*e.g.* in the mouse lung epithelium (Ghosh et al., 2021) or the *Drosophila* gut epithelium (Haller et al., 2017)), or the misexpression of regeneration initiation genes (*e.g*. in the axolotl limb, (Bryant et al., 2017)). In *Platynereis*, the potential effects of serial amputations either on cellular regenerative processes or on the transcriptional responses during regeneration clearly deserve further investigations. Yet, we can hypothesize that the different cellular paths for regeneration depending on the localization, proliferation and differentiation levels of the tissues is important for the morphological robustness of the process.

## Material and methods

### *Platynereis dumerilli* breeding culture, amputation procedure and biological material fixation

*P. dumerilii* juvenile worms were obtained from a husbandry established in the Institut Jacques Monod (for breeding conditions see (Dorresteijn et al., 1993; Vervoort and Gazave, 2022). Standard worms used in experiments were 3-4-month-old with 30-40 segments and were amputated according to the procedure detailed in (Planques et al., 2019; Vervoort and Gazave, 2022). In some experiments, regenerative parts were also re-amputated (either blastemas were totally removed, either regenerative parts were amputated in their middle, see Results section). At given time points reported in the Results section, regenerative parts as well as two posterior-most segments were collected and fixed in 4% paraformaldehyde (PFA) diluted in PBS Tween20 0.1% (PBT) for 2h at RT, then washed in PBT and gradually transferred in 100% Methanol at which point they can be stored at −20°C. For fluorescent beads experiments, the regenerative parts were similarly collected and fixed but were not put in methanol and rather stored up to 24h in PBT at 4°C.

### EdU, BrdU and EdU+BrdU cell proliferation assays

Proliferating cells in S-phase were labelled by incubating the worms with the thymidine analogs EdU (5-Ethynyl-2’-deoxy-Uridine) and/or BrdU (Bromo-deoxy-Uridine), at a respective concentration of 50µM and 1mM in natural fresh sea water (NFSW). Various incubation conditions (duration and biological stage) and pulse and chase experiments were performed as described in the Results section and related figures. The samples were then fixed as described above. Briefly for EdU and/or BrdU labelling, after rehydration, the samples were digested with Proteinase K (40µg/mL for 10min); then the enzyme was inactivated with 2mg/mL Glycine in PBT (1 min) and the samples post-fixed with 4% PFA in PBT (20 min) and washed with PBT. EdU-labelled cells were fluorescently marked by click-it chemistry with the specific addition of a fluorescent azide on EdU molecules (Click-iT™ EdU Cell Proliferation Kit, 488 or 555 dye, ThermoFisher, #C10337 and #C10638), following (Demilly et al., 2013; Vervoort and Gazave, 2022) procedure. BrdU-labelled cells were marked by immunohistochemistry (primary antibody: MoBU-1, mouse, 1:250, ThermoFisher #B35128; secondary antibody: anti-mouse IgG Alexa Fluor® 555 Conjugate, 1:500, goat, Cell Signaling #4409). First, they underwent an antigen retrieval treatment with hydrochloric acid (final concentration of 2M diluted in distilled water for 1h at RT) and were incubated with the antibodies after a blocking step in a solution of sheep serum diluted in PBT. Dual labellings with EdU and BrdU were performed according to a method previously described in (Liboska et al., 2012). The EdU sites were first marked as described above, and the remaining un-labelled EdU sites were then saturated by click-it chemistry with an excess of non-fluorescent azide (Azido-methyl-phenyl-sulfide 95%, Sigma, #244546): the reaction was prepared and achieved according to the manufacturer’s recommendations except the standard fluorescent azide was replaced by the non-fluorescent one at a final concentration of 2mM. The samples were then incubated in hydrochloric acid and the BrdU sites were bound by immunohistochemistry as described previously.

### Probe synthesis and colorimetric whole mount *in situ* hybridization (WMISH)

Probe synthesis and colorimetric whole mount *in situ* hybridization (WMISH) were performed as described in (Demilly et al., 2013; Vervoort and Gazave, 2022). As detailed above for EdU and/or BrdU labelling, after rehydration, the samples were first digested with Proteinase K and post-fixed with PFA. Following that, the samples were pre-hybridized in hybridization buffer at 65 °c (1.5 h), and then incubated with the DIG-labelled probes overnight at 65 °C. DIG was then bound to specific antibodies bearing alkaline phosphatase and the tissues stained thanks to the cleavage of NBT (Nitro Blue Tetrazolium chloride) by this enzyme, catalysed by BCIP (5-Bromo-4-chloro-3-indolyl phosphate). WMISH meant to be observed by bright-field microscopy were mounted in Glycerol. WMISH meant to be observed by confocal microscopy thanks to the reflection procedure (see below) were nuclei counter-stained with Hoechst 0.1% overnight at 4°C and mounted in Glycerol/DABCO (2.5mg/ml DABCO in glycerol). When combined with EdU labelling, the WMISH was first performed just as described. After the coloration with NBT/BCIP, the EdU-labelled cells were fluorescently marked by click-it chemistry as described previously, and then all the nuclei were counter-stained with Hoechst and finally mounted in glycerol/DABCO.

### *In vivo* gut cell labelling with fluorescent beads

To perform *in vivo* labelling of worm gut cells, we used 1µM-diameter fluorescent beads (Fluoresbrite® PolyFluor® 570 Microspheres, Polysciences, #24061-10) that were ingested by the worms. Worms were incubated in a solution of beads diluted in NFSW (1:100) for a week in 24-well plate, one worm per well, in the absence of food. After incubation, worms were rinsed with NFSW to remove non-ingested beads and individually monitored with a fluorescent binocular microscope to determine at which point the gut was marked with fluorescent beads, as the gut is never entirely labelled (the terminal part is always beads-free). Posterior amputations were then performed just anterior to the bead labelling limit, and the worms were let to regenerate in NFSW complemented with food for specific times, depending on the following experiments (see Results section and associated figures). Samples were then collected, counter-stained with Hoechst and mounted in Glycerol/DABCO. When combined with *in vivo* gut cell labelling, EdU labelling was performed by click-it chemistry as seen previously, but for a shorter period of time (5min instead of 1h).

### Cell migration and proliferation inhibitors treatments

Cell proliferation was blocked using Hydroxy-Urea (HU) at 20 mM as previously described (Planques et al., 2019). Cell migration was blocked using LatrunculinB (LatB) at 20nM (as used in (Tweeten and Anderson, 2008)). HU and LatB were dissolved in sea water and DMSO, respectively. Both solutions were changed every 24h to maintain their activities for the duration of the experiment (5 days). Briefly, individual worms were incubated in 2 ml of each solution (or control) on 12-wells plate. LatB-treated worms were scored every day for the regeneration stages that had been reached, as previously described (Planques et al., 2019; Vervoort and Gazave, 2022).

### Histologic samples fixation and sectioning

To perform histological sections, the samples were fixed in a solution of 4% PFA diluted in PBS 1X (without Tween20) for 1h30 at RT and washed in PBS 1X. Then, they were cryoprotected in a solution of sucrose diluted in PBS (300g/L) and stored for 4-5 days at 4°C. After the samples had settled in the sucrose solution, they were transferred into OCT embedding medium (Tissue Freezing Medium, Leica). The remaining sucrose was removed and the samples were put into molds and positioned according to the desired type of section (transverse or longitudinal). The samples were subsequently frozen with dry ice and stored inside the molds at −80°C. The sectioning was performed with a microtome (Leica CM3050S). Sections of 12-14µM were collected on SuperFrost glass slides and stored at −80°C. The cross-sections were then processed for immunostaining (Briscoe et al., 2000), phalloidin labelling (Cytoskeleton, PHDR1; 1/500) or EdU labelling.

### Imaging, acquisition and treatments

Bright-field images of colorimetric WMISH samples were acquired with a Leica CTR 5000 microscope. Fluorescent confocal images of WMISH samples were acquired with a Zeiss LSM780 microscope using a 633nm laser in reflection mode as described in (Jékely and Arendt, 2007). Other fluorescent confocal images were acquired with either a Zeiss LSM780 or LSM980 confocal microscope. Image processing (contrast and brightness, z-projection, auto-blend layers, transversal views) was performed with FIJI and Abode Photoshop. Figures were assembled with Adobe Illustrator.

### EdU+ and/or BrdU+ cell counting

An automatic cell counting procedure was established (see supplementary figure X) and performed with the Imaris software by BitPlane (version 9.5). First, for each sample, all nuclei positions were identified and modelled thanks to the Hoechst signal by using the function “Spots” with a standardized nucleus diameter of 5µm. A Region of Interest (ROI) corresponding specifically, either to the regenerative part or to a body segment was then manually delineated with the surface tool, also thanks to the Hoechst signal and the general morphology of the structure. Then, the spots inside the ROI were sorted along the fluorescent signals of the EdU or/and BrdU with a filter “Intensity Mean” to discriminate true positive nuclei from background. This procedure allowed to determine the absolute number of nuclei inside the ROI and, among them, the number of positive nuclei for each signal; and hence to extract the proportions of EdU+, BrdU+ and EdU+/BrdU+ cells for each sample.

### Statistical analyses

All statistical tests and subsequent graphic representations were performed with GraphPad Prism 7. Mann–Whitney U tests were used to compare whole-blastema proportions of EdU+, BrdU+ and EdU+/BrdU+ cells between different experiments. Wilcoxon signed-rank tests were used to compare those proportions between different parts of the same sample. Multiple Mann–Whitney U tests were used to compare LatB-treated and control worms for each day of treatment. Bonferroni-Dunn test was used afterwards to correct p-values.

## Supplementary figures

**Supplementary Figure 1:**
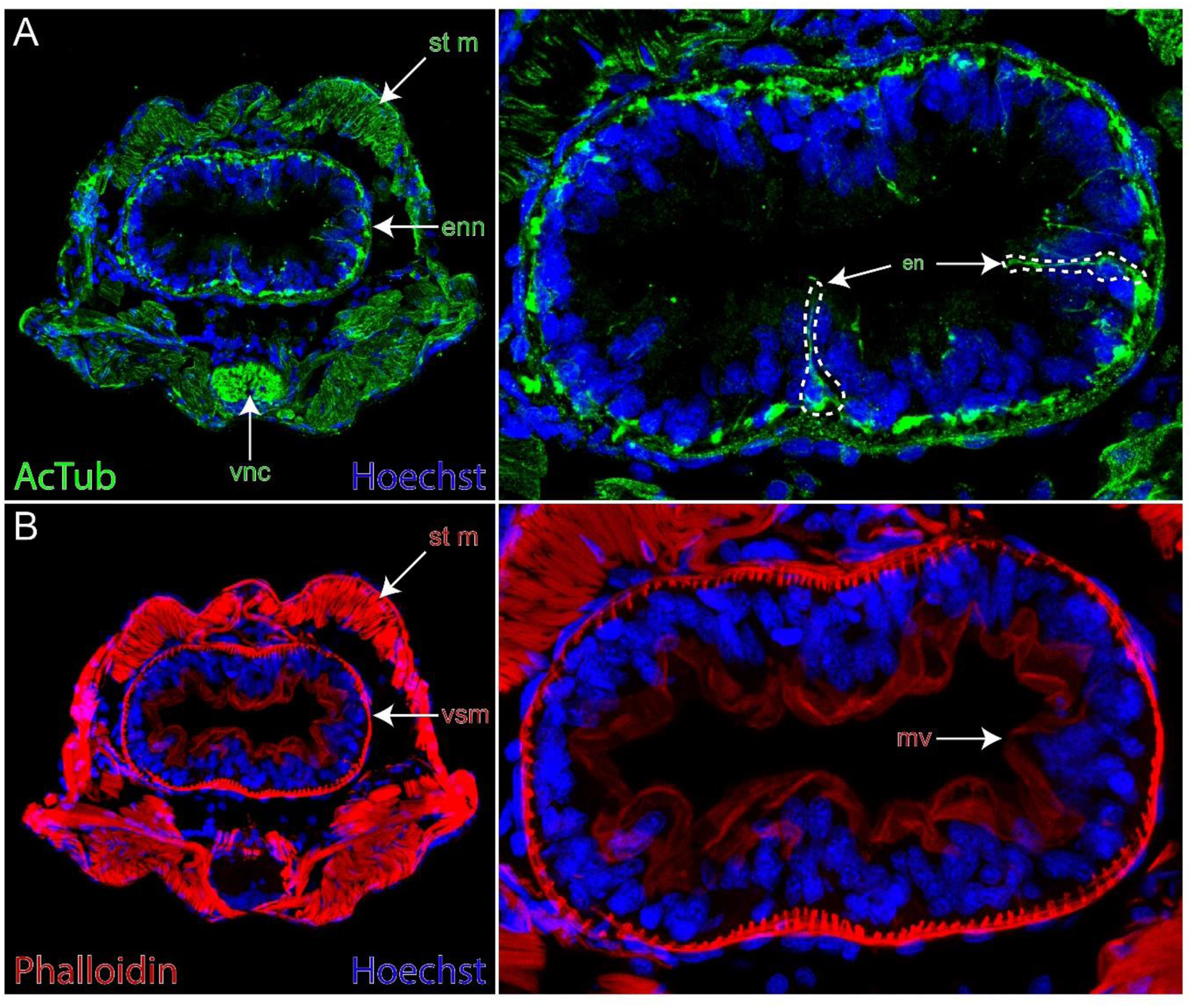
Histological cross-sections on an anterior segment uncover the organization of the gut in *Platynereis*. **(A-B)** Immunolabelling for acetylated tubulin (A, green) or with phalloidin labelling (B, red) and Hoechst DNA staining (blue) performed on a transversal cross-section of an anterior segment. Full transversal cross-sections are shown on the left, while magnifications focused on the gut are shown on the right. st m: striated muscles; enn: enteric nerve net; vnc: ventral nerve cord; en: enteric neuron; vsm: visceral smooth muscles; mv: microvilli.

**Supplementary Figure 2:**
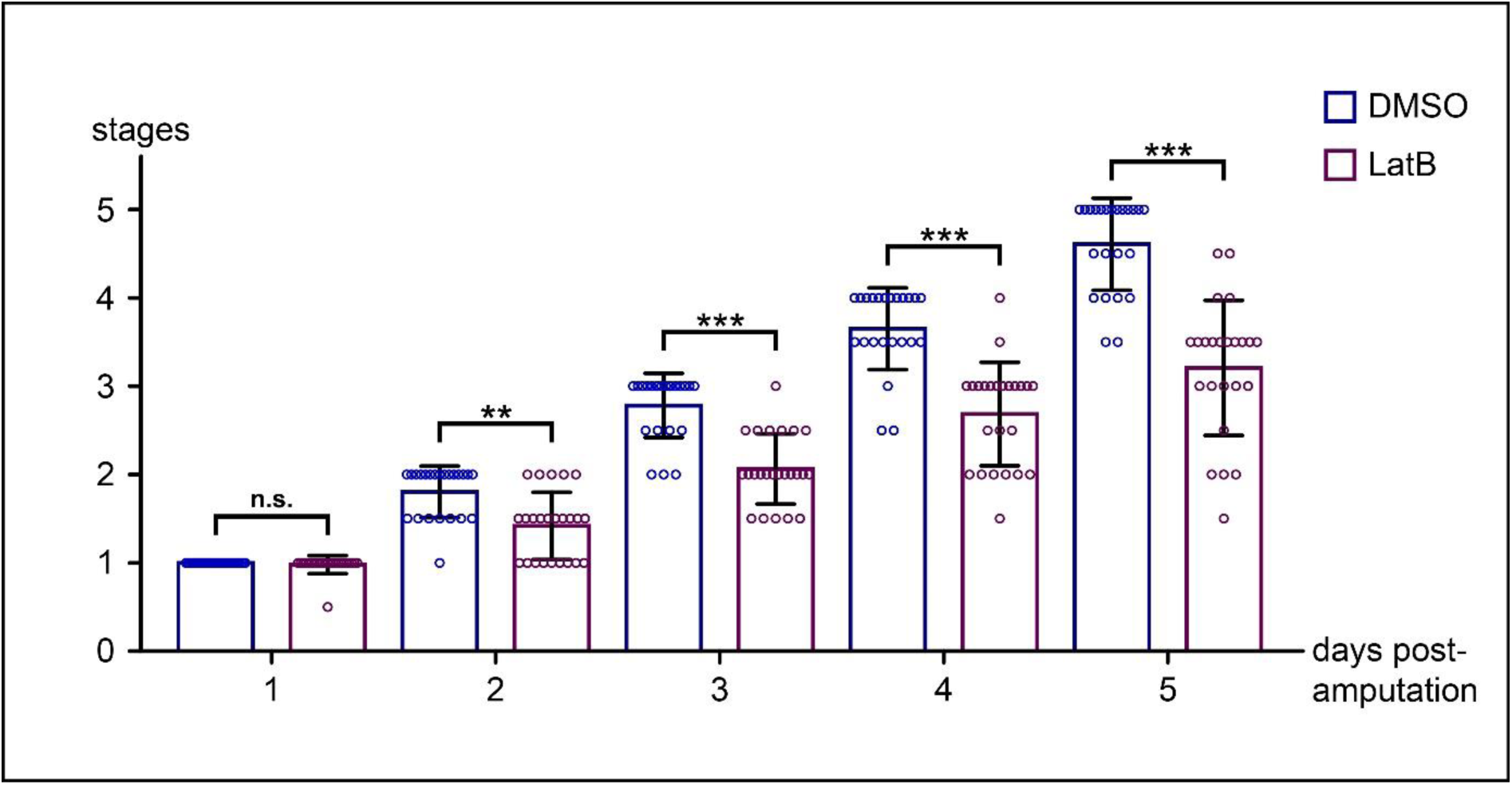
The cell migration inhibitor LatrunculinB impairs posterior regeneration. Graphical representation of the stages reached by control worms (DMSO 0.04 %) and worms treated with 20nM of LatrunculinB or LatB (n=24 for LatB condition and n=23 for control condition) every day during five days. LatB-treated worms showed a significantly delayed regeneration compared to controls from day 2.). Each bracket corresponds to a Mann-Whitney U test. p-values were corrected with Bonferroni-Dunn test. n.s. p >0.05; ** p <0.01; *** p<0.001. Mean + SD are shown.

**Supplementary Figure 3:**
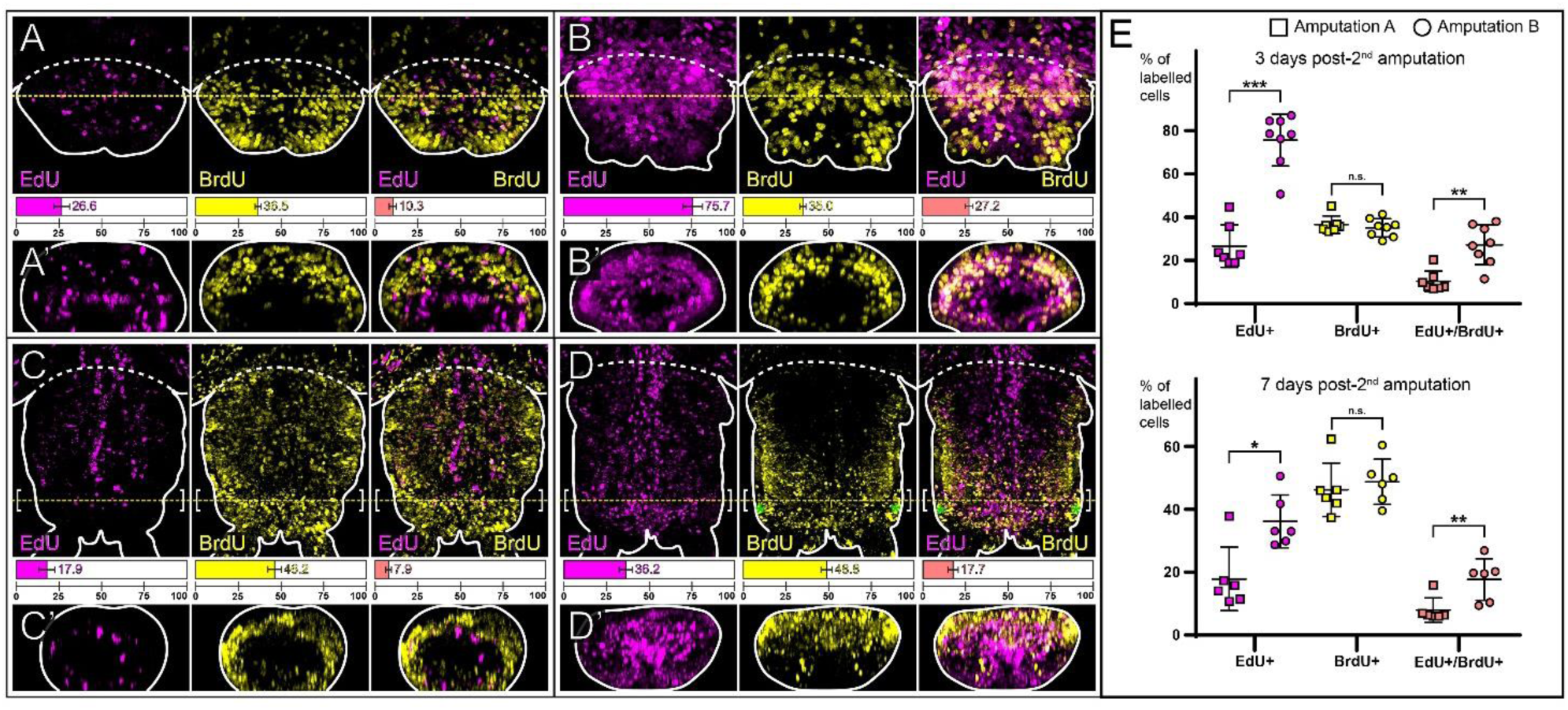
The origin of most of the blastema cells including the stem cells of the growth zone remains local even when regeneration takes place in posterior tissues. Experiments were performed as described in Figure 5A. All worms were amputated posteriorly (amputation “−1”: the growth zone and the pygidium were removed) and let to regenerate for 24h, then incubated with EdU for 5h and let to regenerate for an additional 48h. The worms were then re-amputated: either the blastema and the abutting segment were removed (=amputation A) (A and C), or only the blastema was removed (=amputation B) (B and D). The worms were then let to regenerate for three days, incubated with BrdU (for 1h) and collected (A and B) or let to regenerate for four more days (C and D). **(A-D)** Confocal z-stacks of regenerative parts obtained after both the amputations previously described were performed on worms incubated with EdU (magenta) and BrdU (yellow), (ventral views, anterior is up). **(A’-D’)** Virtual transverse sections along the yellow dotted lines shown in A-D (ventral is up). **(E)** Comparisons of the proportions of EdU+, BrdU+ and EdU+/BrdU+ cells between samples at 3 days post-2^nd^ amputation (A and B) on the top and at 7 days post-2^nd^ amputation (C and D) at the bottom (square = amputation A; circle = amputation B). Each bracket corresponds to a Mann-Whitney U test. n.s. p >0.05; * p <0.05; ** p <0.01; *** p<0.001. Mean + SD are shown. EdU+ and EdU+/BrdU+ cell proportions are significantly higher in B than in A, and in D than in C. In all relevant panels, solid white lines delineate the outlines of the samples, yellow dotted lines indicate the virtual sections plans and white dotted lines correspond to the amputation planes. White brackets indicate the growth zone and green asterisks indicate artefactual staining corresponding to pygidial glands.

**Supplementary Figure 4:**
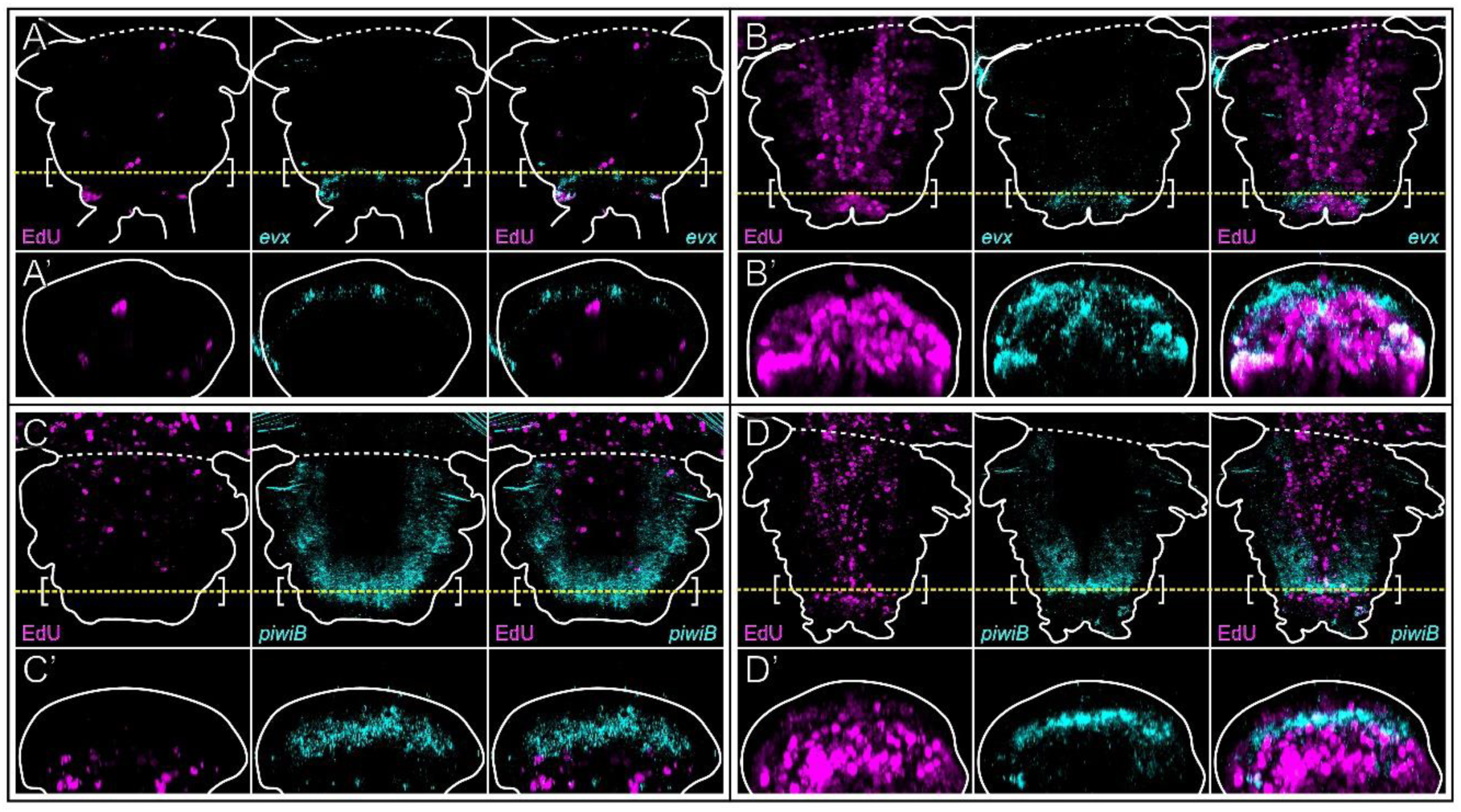
Additional *in situ* hybridizations confirm the local origin of the stem cells of the regenerated growth zone. Experiments were performed as described in Figure 5A. All worms were amputated anteriorly (amputation “−5”: 5 segments and the pygidium were removed) and let to regenerate for 24h, then incubated with EdU for 5h and let to regenerate for an additional 48h. The worms were then re-amputated: either the blastema and the abutting segment were removed (=amputation A) (A and C), or only the blastema was removed (=amputation B) (B and D). The worms were then let to regenerate for seven days. **(A-D)** Confocal z-stacks of regenerative parts at 7 days post-2^nd^ amputation showing EdU+ cells (magenta) as well as *in situ* hybridization signal (cyan) for *evx* (A and B) and *piwiB* (C and D), (ventral views, anterior is up). Corresponding virtual transverse sections (along the yellow dotted lines) are shown in A’ to D’ (ventral is up). EdU+ cells colocalize with *evx* and *piwiB* in B and D but do not in A and C. The solid white lines delineate the outlines of the samples, yellow dotted lines indicate the virtual sections plans and the white dotted lines correspond to the amputation planes. White brackets indicate the growth zone.

## Acknowledgments

This study is dedicated to the memory of our friend, mentor and supervisor, the late Professor Michel Vervoort. We are grateful to all members of the Gazave & Vervoort lab for their support and suggestions on this study. We thank Dr. Quentin Schenkelaars for suggesting the use of fluorescent beads, as well as Dr. Gabriel Krasovec and Dr. Lucie Laplane for helpful comments on the manuscript. We thank Dr. Yves Clément for his help with statistical analyses. We deeply thank the ImagoSeine core facility of Institut Jacques Monod, a member of France-BioImaging (ANR-10-INBS-04) and certified IBiSA.

