## Supplementary figures and images for "Antero-posterior gradients of cell plasticity and proliferation modulate posterior regeneration in the annelid *Platynereis*"

### Supp. Fig. 1

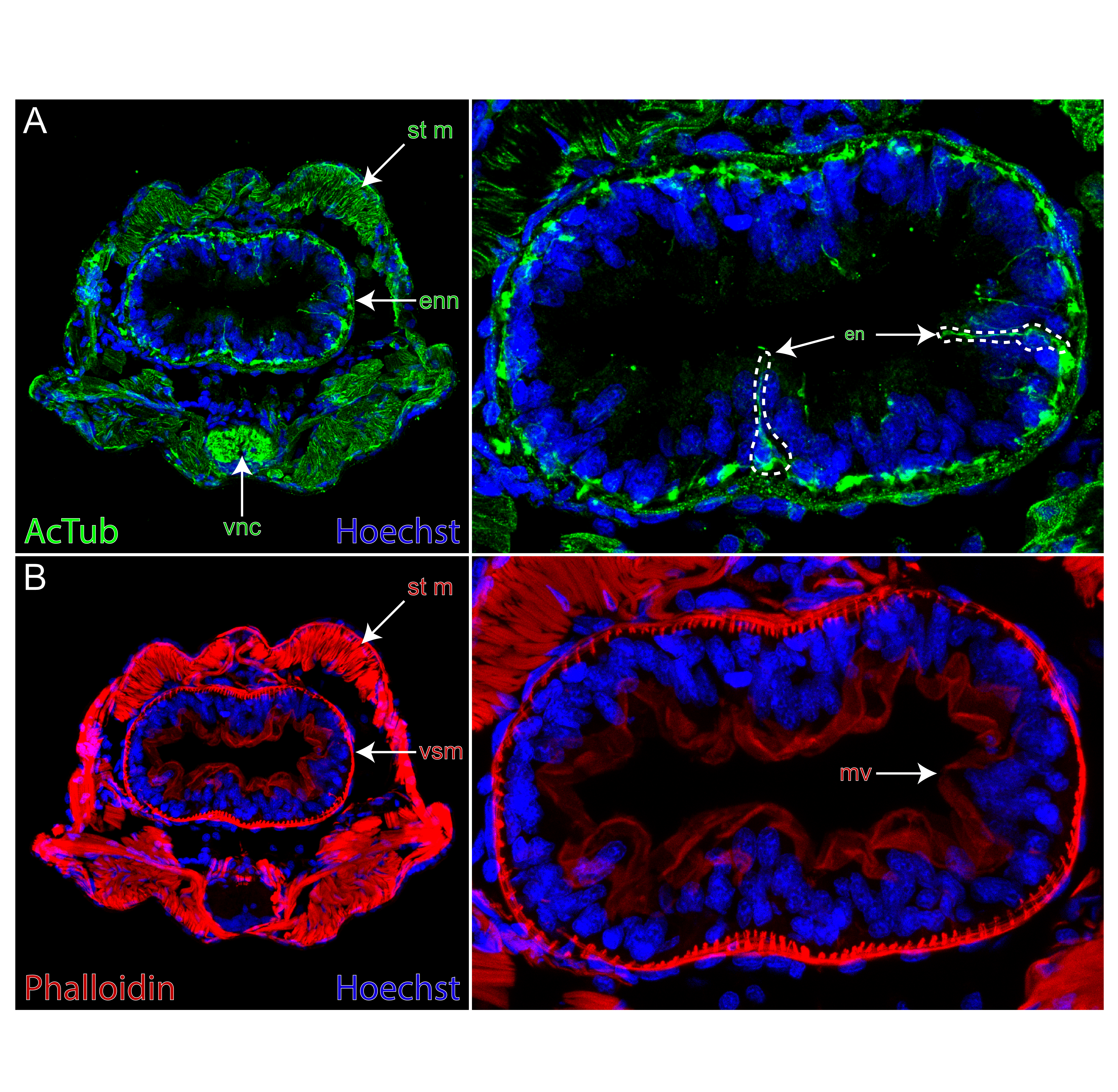

### Supp. Fig. 2

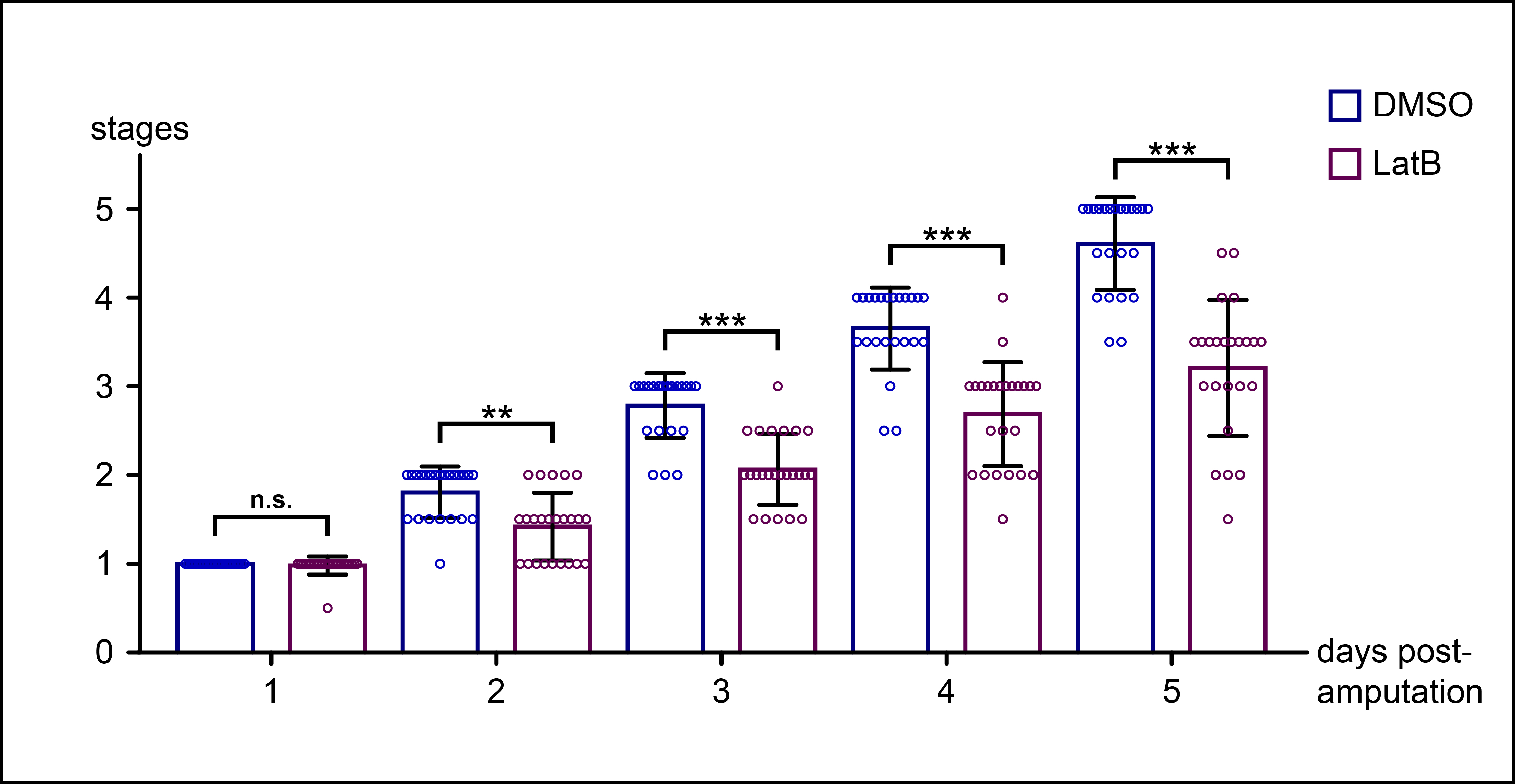

### Supp. Fig. 3

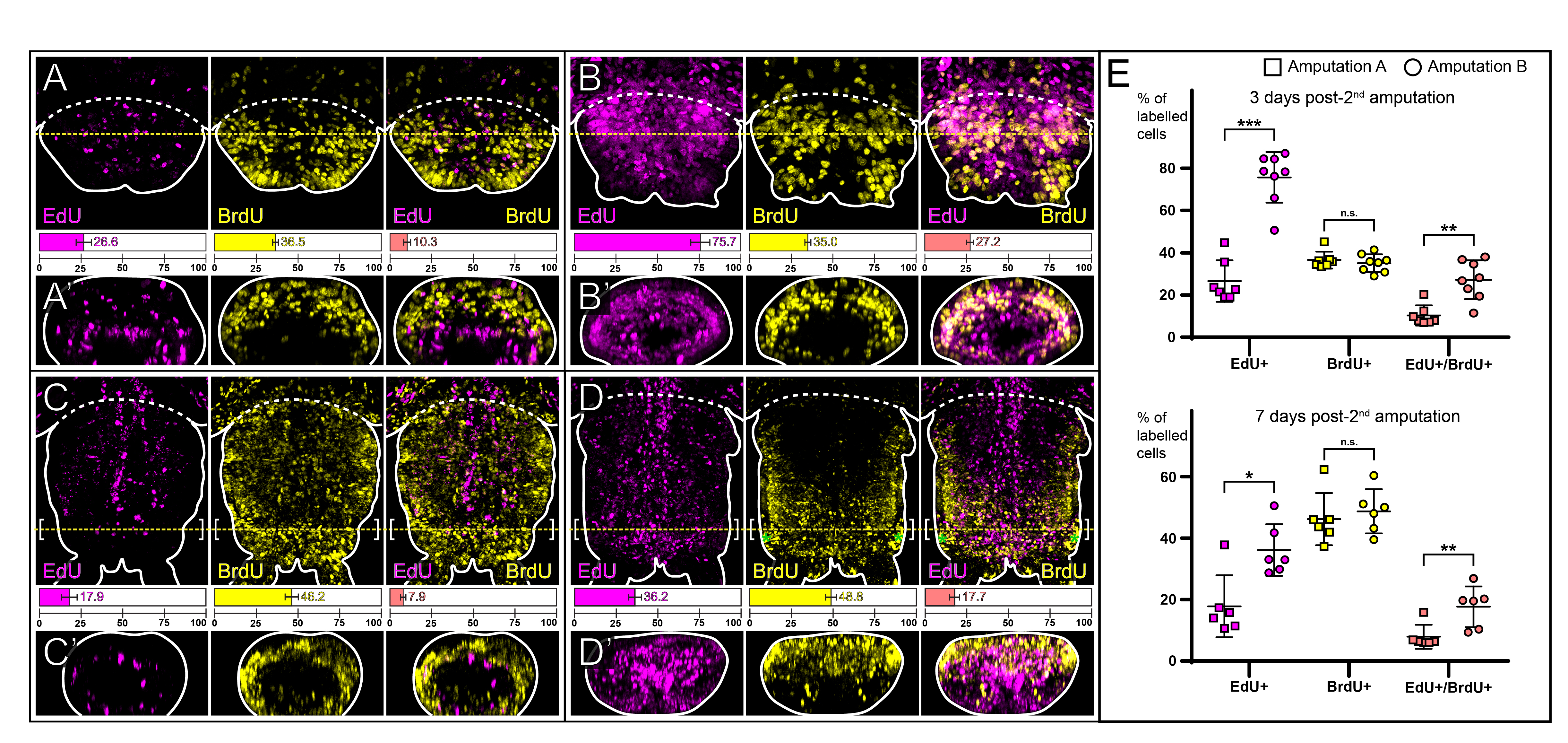

### Supp. Fig. 4

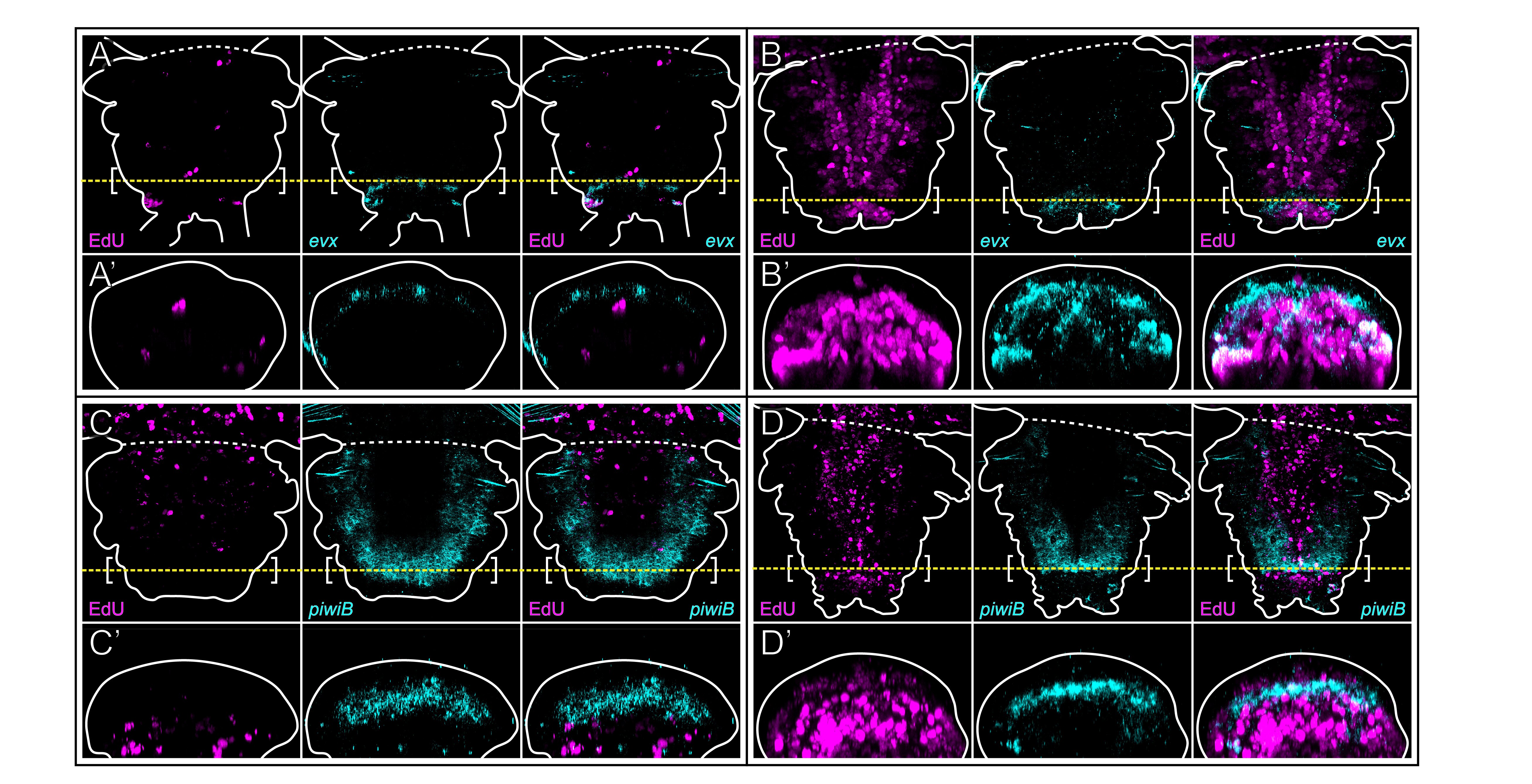
